# Neuronal APOE4 reduction with an APOE-I3-targeting ASO protects against neurodegeneration and neuroinflammation in an Alzheimer’s disease mouse model

**DOI:** 10.64898/2025.12.10.693468

**Authors:** Oscar Yip, Luokang Yao, Jessica Blumenfeld, Min Joo Kim, David Shostak, Zoe Platow, Kaylie Suan, Yaqiao Li, Claire Ellis, Alice An, Yanxia Hao, Qin Xu, Samuel De Leon, Jeremy Nguyen, Yadong Huang

## Abstract

Apolipoprotein E4 (APOE4) is the strongest genetic risk factor for late-onset Alzheimer’s disease (AD). Within the central nervous system (CNS), APOE is produced by a variety of cell types, with differential roles in AD pathogenesis. Studies have shown that APOE4 produced by neurons plays a central role in promoting the development of major AD pathologies, including p-tau accumulation, neuroinflammation, and neurodegeneration, highlighting its role as an upstream initiating factor that affects other cell types and downstream AD-related pathologies. Here, we demonstrate that antisense oligonucleotides (ASOs) targeting APOE-I3, a neuron-specific splicing variant of APOE mRNA, effectively reduce APOE expression in neurons *in vitro* and *in vivo*. Treating PS19 tauopathy mice expressing APOE4 with this APOE-I3-targeting ASO reduces neuronal APOE4, rescues neurodegeneration, and diminishes neuroinflammation. Strikingly, the extent of neuronal APOE4 reduction predicts the efficacy of rescuing neurodegeneration. Single nucleus RNA-sequencing demonstrated that APOE-I3-targeting ASO treatment decreases disease associated neuronal and glial subtypes and increases a disease-protective microglial subtype. These findings suggest that preferential knockdown of neuronal APOE4 with an APOE-I3-targeting ASO protects against key hallmarks of AD pathology, elucidating a potential therapeutic approach for treating APOE4-driven AD.

## INTRODUCTION

Alzheimer’s disease (AD) is a neurodegenerative disorder with pathological changes in the brain that lead to progressive cognitive decline. Despite over a century of AD research, it still lacks effective treatments or cures – likely due to the complex causes of AD pathogenesis, which involves interactions among diverse genetic, epigenetic, and environmental factors. During the past decades, researchers have identified many genetic loci that can alter AD risk with the apolipoprotein E (APOE) locus emerging as the strongest risk factor for late-onset AD^1^.

APOE plays important roles in the brain, such as redistributing lipids among CNS cells for normal lipid homeostasis, repairing injured neurons, maintaining synaptodendritic connections, and scavenging toxins^2^. Within the brain, APOE is expressed primarily by astrocytes, but upon stress, injury, and aging, other cell types, such as neurons and microglia, upregulate their APOE expression^3^. In humans, the *APOE* gene has three common alleles, ε2, ε3 and ε4, and the *APOE* ε4 allele is considered the strongest genetic risk factor, leading to increased AD risk and decreased age of disease onset^1,4,5^. Although numerous studies have demonstrated that APOE4 is linked to more severe AD pathologies^3,6,7^, its roles in AD pathogenesis at a cell type-specific level has been understudied. With more recent studies incorporating single cell omics, it is becoming increasingly clear that the effects of APOE4 on AD pathologies depend on its cellular sources and expression levels^3^.

Previous studies have demonstrated the detrimental effects of APOE4 produced by neurons, including phosphorylated tau (p-tau) accumulation, inhibitory neuron loss, and learning and memory deficits^8–10^. Recently, it has been shown that selective genetic removal of APOE4 from neurons in a tauopathy mouse model resulted in a significant reduction in p-tau pathology, neuroinflammation, and neurodegeneration^11^. Furthermore, it has been reported that neuronal APOE regulates cellular stress and immune response pathways under both health and disease conditions, with the level of neuronal APOE expression tracking with disease progression in patients with MCI and AD^12^. Based on these findings, identifying a strategy to reduce neuronal APOE4 could offer a promising therapeutic approach for APOE4-driven AD.

Antisense oligonucleotides (ASOs) offer an encouraging therapeutic strategy that has the potential to treat neurodegenerative diseases^13,14^, like AD. One key mechanism of ASO action involves sequence-specific binding to target mRNA transcripts, which subsequently recruits the endogenous enzyme RNase H. This recruitment triggers enzymatic cleavage and degradation of the mRNA-ASO duplex, resulting in reduced mRNA stability and ultimately decreased protein translation from the target mRNA^13,15^. For many neurodegenerative diseases that have been linked to the aggregation of toxic proteins, reducing levels of these accumulated proteins can be a powerful therapeutic approach. In fact, ASO drugs are currently in development for the treatment of amyotrophic lateral sclerosis, Huntington’s disease, and AD^16^. When continuously infused intraventricularly, ASOs have been shown to distribute widely throughout the CNS of rodents and primates, including the regions affected in major neurodegenerative diseases^17^. Recent studies treating mice with ASOs via intracerebroventricular (ICV) injection or infusion have found that they can efficiently reduce APOE at both the mRNA and protein levels in the hippocampus and the cortex^18,19^. However, these ASO studies targeted all APOE produced in the brain, and targeting APOE in a cell type-specific manner has not been explored.

Our lab has previously identified a splicing variant of *APOE* mRNA with intron-3 retention, termed *APOE-I3*, which is specific to neurons^20^. This *APOE-I3* transcript was detected in human neuronal cell lines and mouse primary neurons, but was not in human astrocytic cell lines and mouse primary astrocytes^20^. Additionally, RNA-sequencing analysis has shown that the abundance of *APOE-I3* transcripts in the brains of individuals with AD correlates with the severity of tau and amyloid pathologies, and the level of *APOE-I3* transcripts increases with *APOE* ɛ4 allele number^21^. Thus, targeting this neuron-specific APOE4 transcript may be a promising avenue for reducing neuronal APOE4 and its related AD pathologies.

In this study, we designed an ASO targeting intron-3 of the *APOE-I3* transcript and investigated its effects on cell type-specific APOE4 levels and AD pathologies in a human APOE4-expressing tauopathy mouse model. The outcomes of this study provide new insights into how preferential knockdown of neuronal APOE4, using an APOE-I3-targeting ASO, affects the pathogenesis of AD and highlights its promising potential as a therapeutic strategy for treating APOE4-driven AD.

## RESULTS

### Identification of ASO candidates capable of reducing neuronal or global APOE expression

To identify ASOs that target APOE and reduce their expression levels via an RNase-H-mediated knockdown, we designed 20-mer oligonucleotides targeting the *APOE* gene. To preferentially target the neuron-specific splicing variant of *APOE* mRNA with intron-3 retention (APOE-I3 mRNA), we designed five ASOs against the intron-3 sequence of the human *APOE* gene (ASO-I3) (Fig. 1a). Five ASOs against exon regions were also designed to target the *APOE* gene (ASO-E) in any cell type as controls (two targeting exon 3 and three targeting exon 4) (Fig. 1a). To test the efficacy of the designed ASOs, we transfected mouse neuroblastoma Neuro-2a cells stably expressing APOE from a 17-Kb human APOE3 genomic DNA (N2a-E3)^22^ with the ASO candidates.

**Fig. 1.**
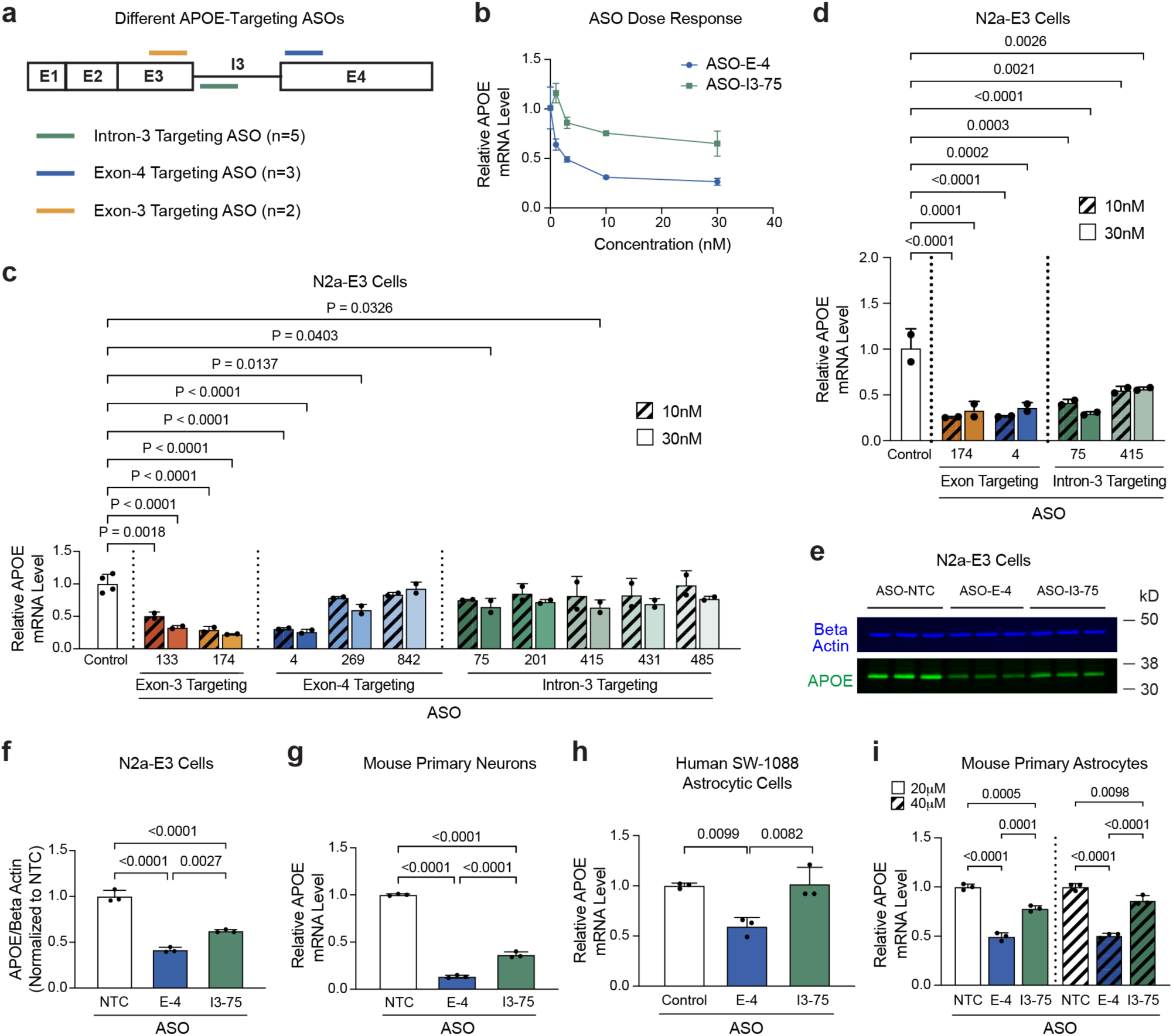
Identification of ASO candidates capable of reducing neuronal or global APOE expression. a, Schematic of APOE-I3 transcript and where differentially designed ASOs target in the intron-3, exon-3, or exon-4 of the APOE-I3 transcript. **b,** qPCR quantification of APOE mRNA levels of neuroblastoma Neuro2a-E3 (N2a-E3) cells treated with an ASO-E-4 or ASO-I3-75 for 24 hours at 0, 1, 3, 10, or 30 nM. **c**, qPCR quantification of APOE mRNA levels of N2a-E3 cells treated with the full panel of 10 ASOs for 24 hours at 10 and 30 nM, respectively. **d**, qPCR quantification of APOE mRNA levels of N2a-E3 cells treated with the four selected ASOs: ASO-E-174, ASO-E-4, ASO-I3-75, and ASO-I3-415 for 48 hours at 10 and 30 nM, respectively. **e**,**f,** Representative western blot images (**e**) and quantification of APOE levels (**f**) in N2a-E3 cells treated with the two selected ASOs: ASO-E-4 and ASO-I3-75, as well as ASO-NTC (a non-targeting control ASO). In **f**, APOE levels were normalized to N2a-E3 cells treated with ASO-NTC. Beta actin was used as loading control. **g**, qPCR quantification of APOE mRNA levels of primary neurons derived from mice expressing human APOE4 from a transgenic APOE4 DNA construct containing intron-3 under the control of a neuron-specific enolase (NSE) promoter (NSE-E4) on a mouse APOE-KO background treated with 10 μM of ASO-NTC, ASO-E-4, or ASO-I3-75 for 1 week. **h**, qPCR quantification of APOE mRNA levels of human SW-1088 astrocytoma cells treated with 3 μM of ASO-NTC, ASO-E-4, or ASO-I3-75 for 48 hours. **i**, qPCR quantification of APOE mRNA levels of primary astrocytes from PS19/E4 mice treated with 20 or 40 μM of ASO-NTC, ASO-E-4, or ASO-I3-75 for 1 week. Throughout, data are expressed as mean ± s.d.. Differences between groups were determined by ordinary one-way ANOVA followed by Dunnett’s multiple comparison test (**c,d**) or ordinary one-way ANOVA followed by Tukey’s multiple comparison test (**f-i**).

To determine the dose-responsive effect of ASOs, we transfected N2a-E3 cells with a representative ASO-I3 and an ASO-E for 24 hours at 0, 1, 3, 10, and 30 nM and measured APOE expression levels via quantitative PCR (qPCR). We found that at 10 nM, ASO-E and ASO-I3 reduced APOE mRNA by ∼69% and ∼25%, respectively (Fig. 1b). At 30 nM, ASO-E and ASO-I3 reduced APOE mRNA by ∼73% and ∼34%, respectively (Fig. 1b). We then decided to treat N2a-E3 cells with the full panel of 10 ASOs for 24 hours at 10 and 30 nM to identify lead ASOs. Two ASO-I3 and two ASO-E (one targeting exon 3 and one targeting exon 4) showed the greatest reduction of APOE mRNA levels (Fig. 1c). The ASO-E-174, targeting exon 3, reduced APOE mRNA by ∼71% and ∼78% at 10 and 30 nM treatment, respectively (Fig. 1c). The ASO-E-4, targeting exon 4, reduced APOE mRNA by ∼69% and ∼74% at 10 and 30 nM treatment, respectively (Fig. 1c). The ASO-I3-75 reduced APOE mRNA by ∼25% and ∼35% at 10 and 30 nM treatment, respectively (Fig. 1c). The ASO-I3-415 reduced APOE mRNA by ∼18% and ∼36% at 10 and 30 nM treatment, respectively (Fig. 1c).

These four ASOs (ASO-E-174, ASO-E-4, ASO-I3-75, and ASO-I3-415) were then selected for further validation and optimization. N2a-E3 were separately treated with the four ASOs for a longer period of 48 hours and the APOE expression levels were measured via qPCR. For 10 and 30 nM treatments, respectively, ASO-E-174 significantly reduced APOE mRNA by ∼78% and ∼67%, ASO-E-4 significantly reduced APOE mRNA by ∼74% and ∼64%, ASO-I3-75 significantly reduced APOE mRNA by ∼59% and ∼71%, and ASO-I3-415 significantly reduced APOE mRNA by ∼45% and ∼44% (Fig. 1d).

Based on their efficacy and region of target in the APOE gene, we selected ASO-E-4 (targeting global APOE expression) and ASO-I3-75 (targeting neuronal APOE expression) for further validation of their effects on APOE protein production. N2a-E3 cells were treated with ASO-E-4 and ASO-I3-75 for 48 hours at 30 nM, and APOE protein levels were determined by western blot analysis relative to cells treated with 30 nM of a non-targeting control ASO (ASO-NTC). Importantly, ASO-E-4 treatment significantly reduced APOE protein levels by ∼58%, and ASO-I3-75 treatment significantly reduced APOE protein levels by ∼38% (Fig. 1e,f). Together, these data demonstrate that both ASO-E-4 and ASO-I3-75 can effectively reduce both APOE mRNA and protein levels in N2a-E3 cells.

### ASO-E-4 and ASO-I3-75 effectively reduce APOE expression in mouse primary neurons

To further examine the efficacy of ASO-E-4 and ASO-I3-75 in neurons, we treated primary neurons derived from mice expressing human APOE4 from a transgenic DNA construct containing intron-3 and under the control of neuron-specific enolase (NSE) promoter (NSE-E4) on a mouse APOE-KO background^23,24^. The primary neuron cultures were treated with 10 μM of ASO-E-4 or ASO-I3-75 on Day 3 for 1 week. Cells were collected on Day 10, and APOE mRNA levels were measured by qPCR. ASO-E-4 significantly reduced APOE mRNA by ∼87% (Fig. 1g), and ASO-I3-75 significantly reduced it by ∼64% (Fig. 1g). These data confirm that both ASO-E-4 and ASO-I3-75 can significantly reduce APOE expression in mouse primary neurons compared to treatment with ASO-NTC.

### ASO-E-4, but not ASO-I3-75, effectively reduces APOE expression in human and mouse astrocytic cells

To test whether the ASO-I3-75 preferentially targets neuronal APOE, we next determined its effect on APOE expression in astrocytes as compared to the ASO-E-4. First, we treated the human astrocytoma cell line, SW-1088, with 3 μM of either ASO for 48 hours and measured APOE mRNA by qPCR. ASO-E-4 significantly reduced APOE mRNA by ∼41% compared to control SW-1088 cells (Fig. 1h). In sharp contrast, ASO-I3-75 did not significantly reduce APOE mRNA, as compared to control SW-1088 cells (Fig. 1h). This suggests that ASO-I3-75 does not effectively alter APOE expression in human astrocytic cells that lack a large, stable pool of APOE-I3 transcripts^20^, whereas ASO-E-4 does effectively reduce APOE expression in human astrocytic cells. These data confirm that ASO-I3-75 effectively reduces APOE expression in neurons without affecting APOE expression in other cell types such as astrocytes.

To further validate the differential efficacy of ASO-I3-75 versus ASO-E-4 in astrocytic cells, we treated primary astrocytes derived from PS19/E4 mice previously generated in our lab^11,25,26^. The cultured astrocytes received either 20 or 40 μM of ASO-E-4, ASO-I3-75, or ASO-NTC on Day 14 for 1 week before qPCR measurement of APOE mRNA levels. ASO-E-4 significantly reduced APOE mRNA by ∼51% and ∼49% at the 20 and 40 μM treatments, respectively, while ASO-I3-75 only reduced APOE mRNA by ∼22% and ∼14% at the same doses (Fig. 1i). Taken together, these data demonstrate that in primary mouse astrocytes, ASO-E-4 effectively reduces APOE expression (∼50%), while ASO-I3-75 does not effectively do so (∼15–20%), thus highlighting ASO-I3-75 as a neuron-preferential APOE ASO.

### ASO-E-4 and ASO-I3-75 effectively reduce APOE expression in the hippocampus of PS19/E4 mice in a cell-type-dependent manner

To determine the efficacy of ASO-E-4 and ASO-I3-75 in reducing APOE expression in an AD mouse model, we next treated PS19/E4 tauopathy mice previously generated in our lab^11,25,26^ with the ASOs. To best characterize the therapeutic potential of APOE reduction, we began ASO treatment at 8.5 months of age, when mice have already begun developing some AD pathologies. Treatment was conducted via an osmotic pump ICV infusion, allowing for slow delivery of 850 ug of ASO-E-4, ASO-I3-75, or ASO-NTC for 4 weeks (Fig. 2a). Following treatment, APOE protein levels were quantified via immunostaining for APOE and determining APOE percent coverage area in the dentate gyrus (DG) and CA1 hippocampal subregions. PS19/E4 mice treated with either ASO-E-4 or ASO-I3-75 had a significant reduction in overall APOE protein levels, as compared to PS19/E4 mice treated with ASO-NTC (Fig. 2b,c). Thus, treatment of PS19/E4 mice with ASO-E-4 or ASO-I3-75 via osmotic pump infusion into the ventricle for 4 weeks effectively reduces APOE protein levels in the hippocampus.

**Fig. 2.**
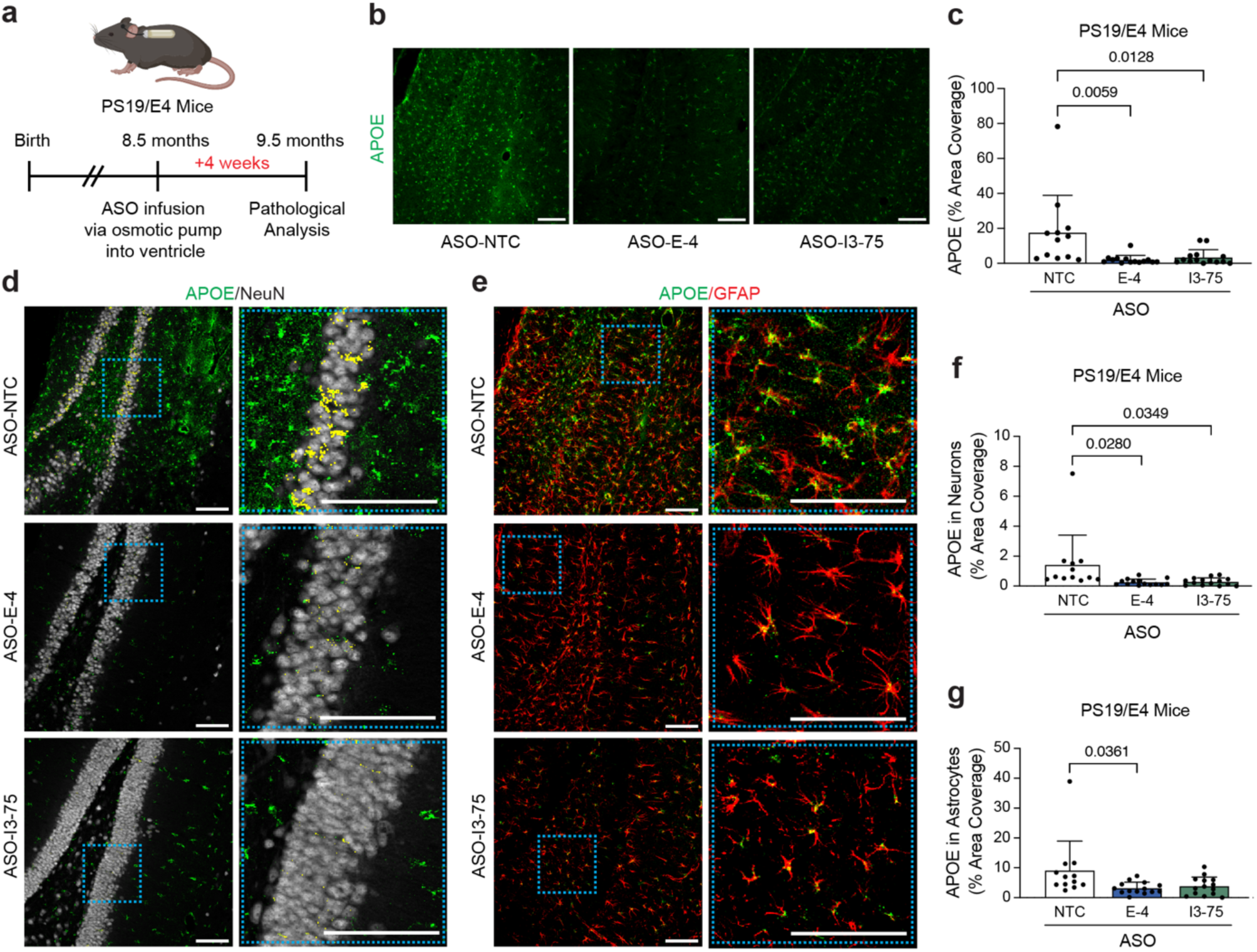
ASO-E-4 and ASO-I3-75 effectively reduce APOE expression in the hippocampus of PS19/E4 mice in a cell-type-dependent manner. a, Schematic timeline of ASO treatment via osmotic pump infusion into ventricle and pathological analysis of PS19/E4 mice. **b**, Representative images showing immunostaining of APOE in the hippocampus of PS19/E4 mice treated with ASO-NTC, ASO-E-4, or ASO-I3-75 for four weeks. Scale bars are 100μm. **c**, Quantification of APOE percent area coverage in the dentate gyrus (DG) and CA1 of PS19/E4 mice treated with ASO-NTC, ASO-E-4, or ASO-I3-75. For all mice, data is averaged from 4 fields of view from 2 brain sections (2 fields of view from the DG and 2 fields of view from the CA1). ASO-NTC, n=12 mice; ASO-E-4, n=14 mice; ASO-I3-75, n=14 mice. **d**, Representative images showing immunostaining of APOE (green) and NeuN (white) in the DG of PS19/E4 mice treated with ASO-NTC, ASO-E-4, or ASO-I3-75. Each representative image includes a magnified image of a region in the DG to the right, with areas positive for both APOE and NeuN pseudo-colored in yellow for clear visualization. Scale bars are 100μm. **e**, Representative images showing immunostaining of APOE (green) and GFAP (red) in the DG of PS19/E4 mice treated with ASO-NTC, ASO-E-4, or ASO-I3-75. Each representative image includes a magnified image of a region in the DG to the right, with areas positive for both APOE and GFAP in yellow. Scale bars are 100μm. **f,** Quantification of APOE percent area coverage in neurons of PS19/E4 mice treated with ASO-NTC, ASO-E-4, or ASO-I3-75. **g**, Quantification of APOE percent area coverage in astrocytes of PS19-E4 mice treated with ASO-NTC, ASO-E-4, or ASO-I3-75. Throughout, data are expressed as mean ± s.d.. Differences between groups were determined by ordinary one-way ANOVA followed by Tukey’s multiple comparison test.

To gain a better understanding of cell type-specific APOE reduction from the two ASOs, we measured the APOE percent coverage area in neurons, labeled with NeuN, and astrocytes, labeled with GFAP. PS19/E4 mice treated with either ASO-E-4 or ASO-I3-75 showed a significant reduction in neuronal APOE levels compared to PS19/E4 mice treated with ASO-NTC (Fig. 2d,f). Meanwhile, for astrocytic APOE levels, PS19/E4 mice treated with ASO-E-4 exhibited a significant reduction relative to mice treated with ASO-NTC, while PS19/E4 mice treated with ASO-I3-75 showed a trending but not significant reduction of APOE (Fig. 2e,g). Thus, ASO-E-4 and ASO-I3-75 can effectively reduce APOE expression in the hippocampus of PS19/E4 mice in a cell-type-dependent manner.

### ASO-I3-75, but not ASO-E-4, effectively reduces neurodegeneration in the hippocampus of PS19/E4 mice

Next, we evaluated the effects of the two ASOs on neurodegeneration, as determined by hippocampal volume, in the same PS19/E4 mice after 4 weeks of ASO treatment (at 9.5 months of age). Hippocampal volume was significantly higher in PS19/E4 mice treated with ASO-I3-75 compared to those treated with ASO-NTC (Fig. 3a,b), showing reduced hippocampal atrophy. Interestingly, PS19/E4 mice treated with ASO-E-4 exhibited a trending, but not significant, increase in hippocampal volume relative to those treated with ASO-NTC (Fig. 3a,b). These findings suggest that preferentially reducing neuronal APOE4 by targeting the neuron-specific splicing variant APOE-I3, as ASO-I3-75 did, is more efficient and beneficial for rescuing neurodegeneration, as compared to targeting global APOE4 in the brain, as ASO-E-4 did.

**Fig. 3.**
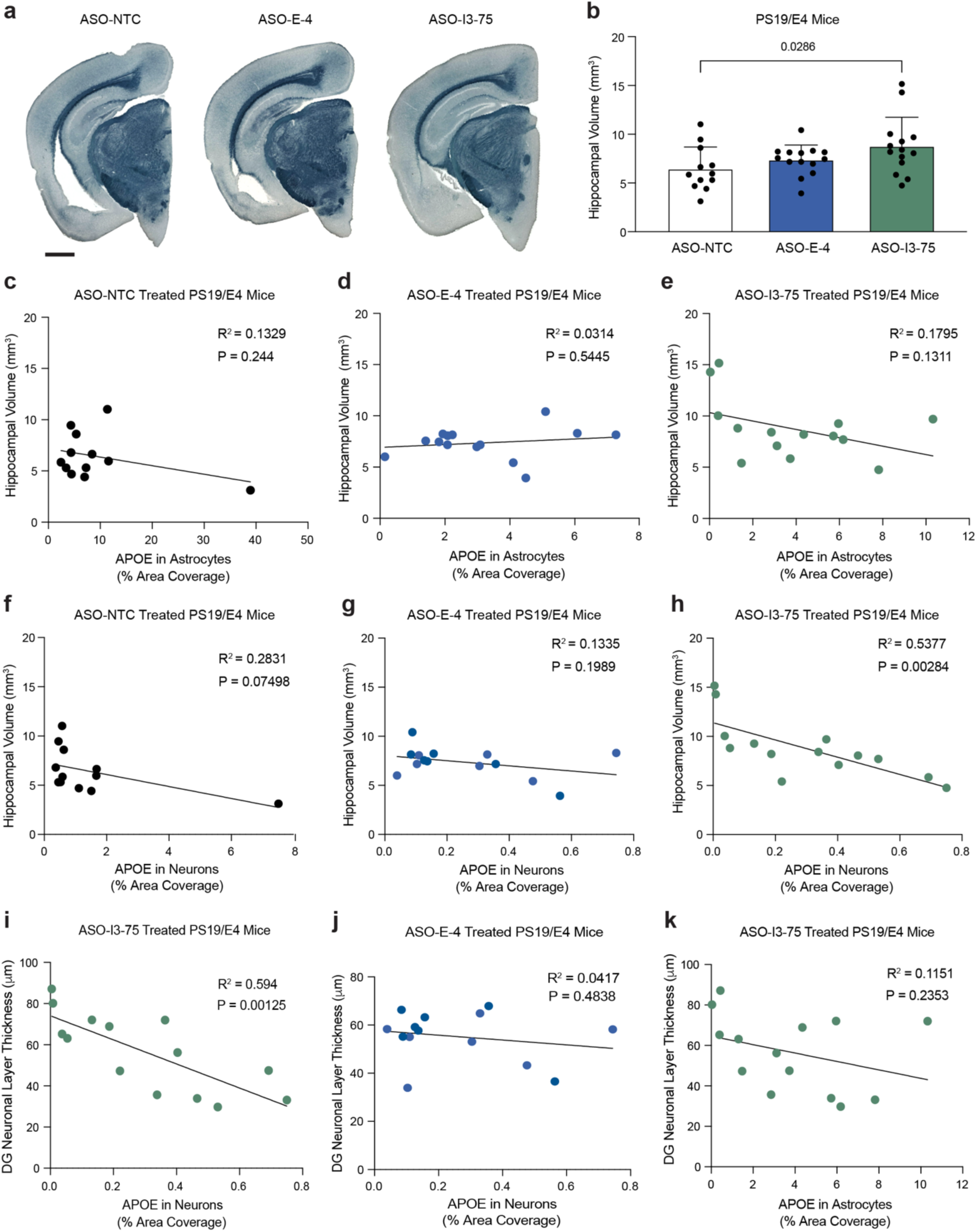
ASO-I3-75, but not ASO-E-4, effectively reduces neurodegeneration in the hippocampus of PS19/E4 mice. a, Representative images of the ventral hippocampus of 9.5-month-old PS19/E4 mice treated with ASO-NTC, ASO-E-4, or ASO-I3-75 for 4 weeks after staining with Sudan Black to enhance hippocampal visualization. ASO-NTC, n=12 mice; ASO-E-4, n=14 mice; ASO-I3-75, n=14 mice. Scale bar is 1mm. **b**, Quantification of hippocampal volume in 9.5-month-old PS19/E4 mice treated with ASO-NTC, ASO-E-4, or ASO-I3-75 for 4 weeks. Data are expressed as mean ± s.d., and differences between groups were determined by ordinary one-way ANOVA followed by Dunnett’s multiple comparison test**. c-e**, Correlations between hippocampal volume (mm^3^) and APOE level in astrocytes (% area coverage) in PS19/E4 mice treated with ASO-NTC (**c,** n=12), ASO-E-4 (**d**, n=14), or ASO-I3-75 (**e**, n=14). **f-h**, Correlations between hippocampal volume (mm^3^) and APOE level in neurons (% area coverage) in PS19/E4 mice treated with ASO-NTC (**f**), ASO-E-4 (**g**), or ASO-I3-75 (**h**). **i**, Correlations between DG neuronal layer thickness (μm) and APOE level in neurons (% area coverage) in PS19/E4 mice treated with ASO-I3-75 (n=14). **j**, Correlations between DG neuronal layer thickness (μm) and APOE level in neurons (% area coverage) in PS19/E4 mice treated with ASO-E-4 (n=14). **k**, Correlations between DG neuronal layer thickness (μm) and APOE level in astrocytes (% area coverage) in PS19/E4 mice treated with ASO-I3-75 (n=14). For all correlations, Pearson’s correlation analysis (two-sided) was performed.

To gain a better understanding of how APOE4 levels in different cell types contribute to neurodegeneration, we assessed the relationship between hippocampal volume and APOE4 levels in neurons and astrocytes. Strikingly, there were no significant correlations between astrocytic APOE4 levels and hippocampal volumes in mice treated with either ASO-NTC, ASO-E-4, or ASO-I3-75 (Fig. 3c-e), suggesting that astrocytic APOE4 is not a driver of APOE4-promoted hippocampal degeneration in PS19/E4 mice. In sharp contrast, there was a strong negative correlation between neuronal APOE4 levels and hippocampal volumes in PS19/E4 mice treated with ASO-I3-75 (Fig. 3h), although such a correlation was not present in the mice treated with either ASO-NTC or ASO-E-4 (Fig. 3f,g). Thus, neuron-preferential reduction of APOE4 likely drives the protection of hippocampal degeneration.

In line with these observations, there was also a strong negative correlation between neuronal APOE4 levels and DG NeuN+ granular layer thickness, another measurement of neurodegeneration^11,25^, in PS19/E4 mice treated with ASO-I3-75 (Fig. 3i), but not in those treated with ASO-E-4 (Fig. 3j). Importantly, there was no significant correlation between astrocytic APOE4 levels and DG NeuN^+^ granular layer thickness in PS19/E4 mice treated with either ASO-I3-75 (Fig. 3k) or ASO-E-4 (Extended Data Fig. 1a). There was a trending, but not statistically significant, increase in the average DG NeuN^+^ granular layer thickness in PS19/E4 mice treated with either ASO-I3-75 or ASO-E-4 versus those treated with ASO-NTC (Extended Data Fig. 1b). Notably, for mice treated with ASO-NTC, ASO-E-4, or ASO-I3-75, DG NeuN+ granular layer thickness had a significant positive correlation with hippocampal volume, suggesting that neuronal loss in the DG contributes to the observed hippocampal volume reduction (Extended Data Fig. 1c-e). Together, these findings suggest that preferentially reducing neuronal APOE4 with ASO-I3-75 treatment leads to efficient rescue of neurodegeneration, including hippocampal atrophy and DG neuronal layer shrinkage.

### ASO-I3-75, but not ASO-E-4, diminishes p-tau contribution to neurodegeneration in the hippocampus of PS19/E4 mice

To determine whether treatment with the different APOE-targeting ASOs protects against p-tau accumulation in PS19/E4, we immunostained brain sections with a p-tau-specific AT8 monoclonal antibody^11,25,27^. There were no significant differences in AT8^+^ percent area coverage in the hippocampus between mice treated with ASO-E4, ASO-I3-75, or ASO-NTC (Extended Data Fig. 2a-d). Correlation analysis revealed that p-tau (AT8^+^) percent coverage areas negatively correlated with hippocampal volumes in PS19/E4 mice treated with ASO-NTC (Extended Data Fig. 2e), suggesting the contribution of p-tau to neurodegeneration. Strikingly, although such a negative correlation was still present in PS19/E4 mice treated with ASO-E-4 (Extended Data Fig. 2f), it was no longer present in the mice treated with the neuronal-APOE-targeting ASO-I3-75 (Extended Data Fig. 2g). Similarly, there was a negative correlation between p-tau (AT8^+^) coverage area and DG NeuN^+^ layer thickness in PS19/E4 mice treated with ASO-NTC or ASO-E-4, which was also no longer present in the PS19/E4 mice treated with ASO-I3-75 (Extended Data Fig. 2h-j). Taken together, all these data indicate that neuron-preferential reduction of APOE4 diminishes p-tau contribution to neurodegeneration in the hippocampus of PS19/E4 mice, and this protective effect does not apply to global APOE4 reduction.

### ASO-E-4 and ASO-I3-75 reduces astrogliosis in the hippocampus of PS19/E4 mice

We then tested whether treatment with the different APOE-targeting ASOs protects against APOE4-driven gliosis in the same cohort of PS19/E4 mice. To assess for astrogliosis, we stained with a GFAP-specific antibody (Fig. 4a) and calculated for GFAP^+^ percent coverage area in the hippocampus^11,25^. PS19/E4 mice treated with either ASO-E-4 or ASO-I3-75 exhibited a significantly lower percent coverage area of GFAP^+^ astrocytes in the DG and CA1 of the hippocampus relative to mice treated with ASO-NTC (Fig. 4a,b), indicating that both global and neuron-preferential reduction of APOE4 decreases astrogliosis. There was a significant negative correlation between GFAP^+^ percent coverage area and hippocampal volume across all treatment groups (Fig. 4i), suggesting that in PS19/E4 mice, astrogliosis is a good indicator (or potentially a contributing factor) to hippocampal degeneration.

**Fig. 4.**
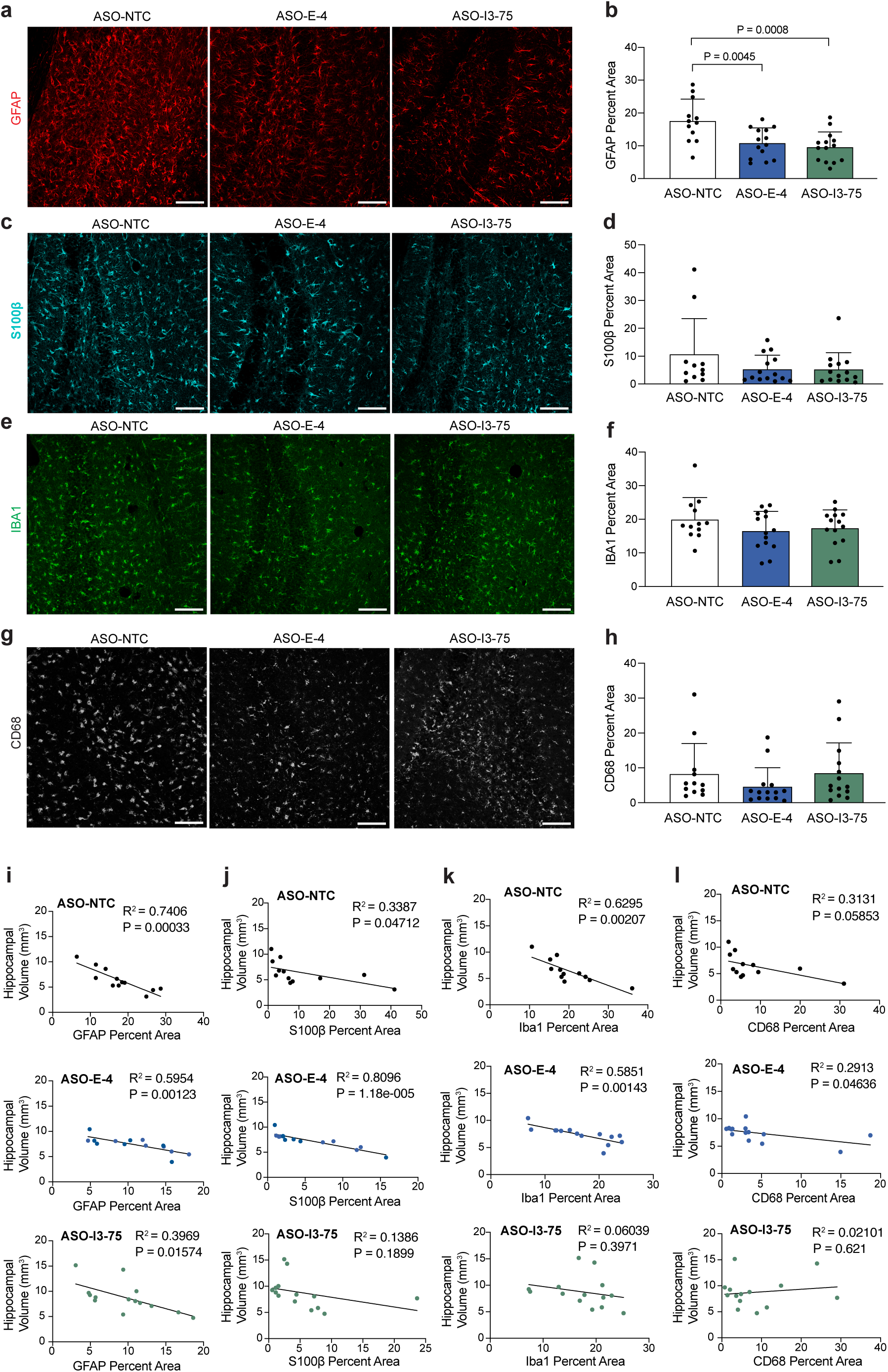
Both ASO-E-4 and ASO-I3-75 reduces astrogliosis while only ASO-I3-75 alters the contribution of microgliosis to hippocampal degeneration in PS19/E4 mice. a, Representative images of astrocyte immunostaining with anti-GFAP in the dentate gyrus (DG) of 9.5-month-old PS19/E4 mice treated with ASO-NTC, ASO-E-4, or ASO-I3-75 for 4 weeks. **b**, Quantification of percent GFAP coverage area in the DG and CA1 of 9.5-month-old PS19/E4 mice treated with ASO-NTC, ASO-E-4, or ASO-I3-75. **c,** Representative images of activated astrocyte immunostaining with anti-S100β in the DG of 9.5-month-old PS19/E4 mice treated with different ASOs. **d**, Quantification of percent S100β coverage area in the DG and CA1 of 9.5-month-old PS19/E4 mice treated with different ASOs. **e**, Representative images of microglia immunostaining with anti-Iba1 in the DG of 9.5-month-old PS19/E4 mice treated with different ASOs. **f**, Quantification of the percent Iba1 coverage area in the DG and CA1 of 9.5-month-old PS19/E4 mice treated with different ASOs. **g**, Representative images of activated microglia immunostaining with anti-CD68 in the DG of 9.5-month-old PS19/E4 mice treated with different ASOs. **h**, Quantification of the percent CD68 coverage area in the DG and CA1 of 9.5-month-old PS19/E4 mice treated with different ASOs. For all images, scale bars are 100μm. For all image quantifications, data is averaged from 4 fields of view from 2 brain sections (2 fields of view from the DG and 2 fields of view from the CA1). ASO-NTC, n=12 mice; ASOE-4, n=14 mice; ASO-I3-75, n=14 mice. **i**, Correlations between hippocampal volume (mm^3^) and percent GFAP coverage area in PS19/E4 mice treated with ASO-NTC, ASO-E-4, or ASO-I3-75. **j**, Correlations between hippocampal volume (mm^3^) and percent S100β coverage area in PS19/E4 mice treated with ASO-NTC, ASO-E-4, or ASO-I3-75. **k**, Correlations between hippocampal volume (mm^3^) and percent IBA1 coverage area in PS19/E4 mice treated with ASO-NTC, ASO-E-4, or ASO-I3-75. **l**, Correlations between hippocampal volume (mm^3^) and percent CD68 coverage area in PS19/E4 mice treated with ASO-NTC, ASO-E-4, or ASO-I3-75. For all quantifications of percent area coverage, data are expressed as mean ± s.d., and differences between groups were determined by ordinary one-way ANOVA followed by Tukey’s multiple comparison test. For all correlations, Pearson’s correlation analysis (two-sided) was performed.

We also assessed the percent coverage area of activated astrocytes by staining for S100β (Fig. 4c). PS19/E4 mice treated with either ASO-E-4 or ASO-I3-75 exhibited a trending, but not significant reduction of percent coverage area of S100β^+^ astrocytes relative to mice treated with ASO-NTC (Fig. 4c,d). However, when correlating S100β^+^ percent coverage area with hippocampal volume, there was a significant negative correlation in PS19/E4 mice treated with either ASO-NTC or ASO-E-4 (Fig. 4j). Interestingly, there was no longer a significant negative correlation in PS19/E4 mice treated with ASO-I3-75 (Fig. 4j). Collectively, these data suggest that preferentially reducing neuronal APOE4 eliminates the contribution of activated astrocytes to hippocampal degeneration.

### ASO-I3-75, but not ASO-E-4, alters the contribution of microgliosis to hippocampal degeneration in PS19/E4 mice

Next, to assess the effects of different APOE-targeting ASOs on microgliosis in the same cohort of PS19/E4 mice, we immunostained for IBA1 and analyzed the IBA1^+^ percent coverage area in the hippocampus (Fig. 4e). PS19/E4 mice treated with either ASO-E-4 or ASO-I3-75 did not exhibit a significant (although trending) reduction of the percent coverage area of IBA1^+^ microglia, as compared to the mice treated with ASO-NTC (Fig. 4e,f). When correlating IBA1^+^ percent coverage area to hippocampal volume, similar to the case with S100β, there was a significant negative correlation in PS19/E4 mice treated with either ASO-NTC or ASO-E-4, but not in mice treated with ASO-I3-75 (Fig. 4k). These data suggest that microgliosis is a good indicator or contributing factor to hippocampal degeneration, as seen in PS19/E4 mice treated with ASO-NTC or ASO-E4-4; however, neuron-preferential reduction of APOE4 eliminates such a relationship.

We also assessed the percent coverage of activated microglia by staining for CD68 in the same cohort of mice (Fig. 4g). PS19/E4 mice treated with either ASO-E-4 or ASO-I3-75 did not exhibit a significant change in the percent coverage area of CD68^+^ microglia compared to the mice treated with ASO-NTC (Fig. 4g,h). When correlating CD68^+^ percent coverage area to hippocampal volume, there was a strong trending negative correlation in PS19/E4 mice treated with ASO-NTC and a significant negative correlation in PS19/E4 mice treated with ASO-E-4 (Fig. 4l). However, there was no longer a significant negative correlation in PS19/E4 mice treated with ASO-I3-75. Again, these data suggest that increased activated microglia is a good indicator or contributing factor to hippocampal degeneration, as seen in PS19/E4 mice treated with ASO-NTC or ASO-E4-4; however, neuron-preferential reduction of APOE4 eliminates such a relationship. In other words, in mice treated with ASO-I3-75, activated microglia are no longer a good indicator of hippocampal degeneration, perhaps due to shifts in disease-associated and disease-protective subpopulations of microglia in response to decreased neuronal APOE4 expression (see single nucleus transcriptomic data in later sections). Taken together, the contributions of microglia or activated microglia to hippocampal degeneration depend on neuronal expression of APOE4.

### ASO-E-4 and ASO-I3-75 diminishes disease-associated excitatory neurons in the hippocampus of PS19/E4 mice

To gain an in-depth understanding of the cell type-specific effects of the two APOE-targeting ASOs at a transcriptomic level, we performed single nucleus RNA-sequencing (snRNA-seq) on isolated hippocampi from the same cohort of PS19/E4 mice after 4 weeks of ASO treatment (at 9.5 months of age). The snRNA-seq dataset contained 161,796 nuclei covering 26,885 genes after normalization and filtering for quality control. Clustering by the Louvain algorithm and visualization by Uniform Manifold Approximation and Projection (UMAP) revealed 38 distinct cell clusters (Fig. 5a). Based on their expression of marker genes (Supplementary Table 1), these clusters were assigned to 13 excitatory neuron (Ex) clusters (2-4, 7, 9, 16, 22, 27, 28, 30, 31, 34, and 37), 4 inhibitory neuron (In) clusters (8, 11, 15, and 26), 7 subiculum clusters (10, 14, 17, 18, 20, 25, and 33), 3 oligodendrocyte clusters (1, 5, and 24), 1 astrocyte cluster (13), 2 microglia clusters (6 and 21), 3 OPC clusters (12, 29, and 38) and 4 unknown clusters (23, 32, 35, and 36) (Fig. 5a and Extended Data Fig. 3a,b).

**Fig. 5.**
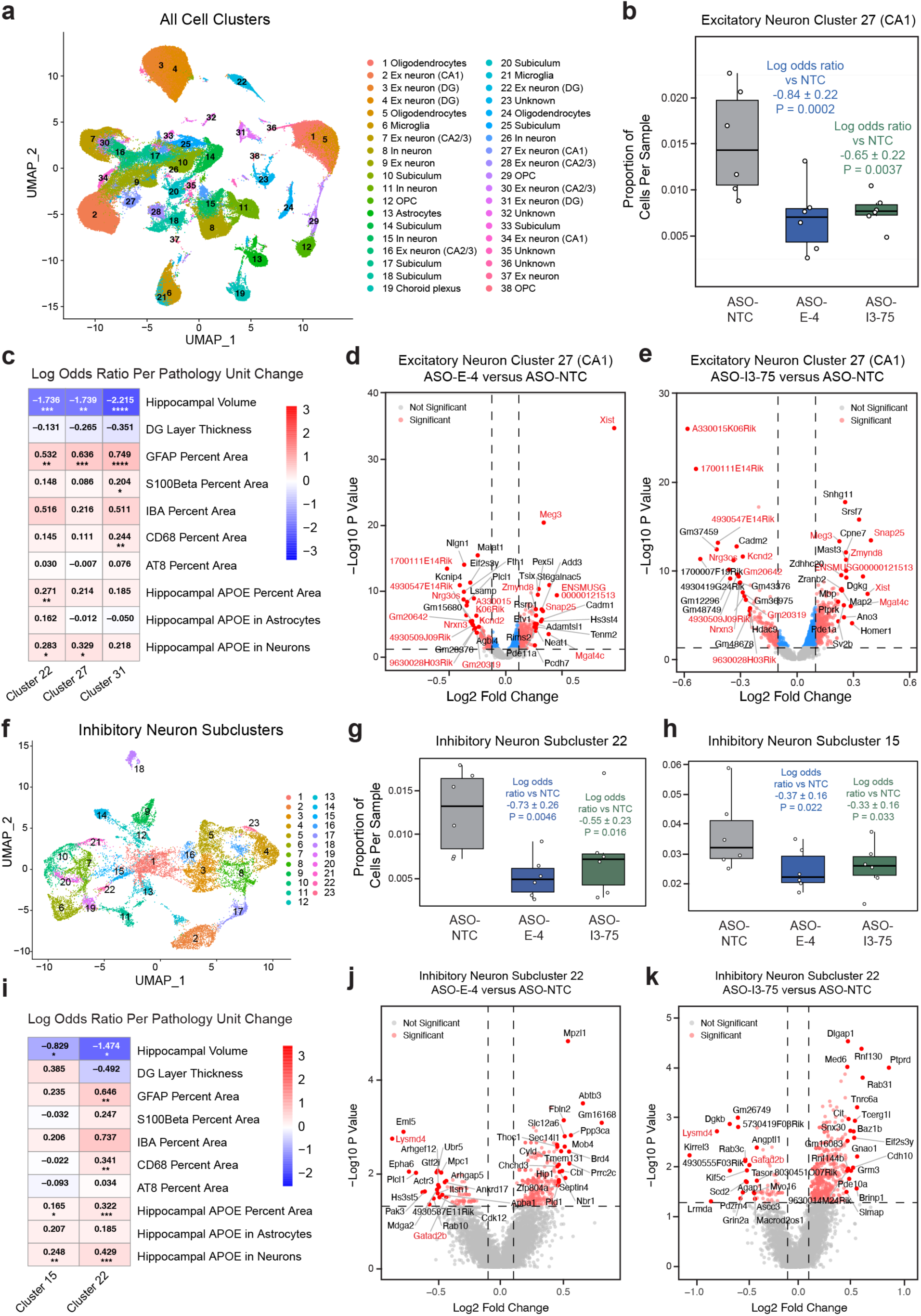
ASO-E-4 and ASO-I3-75 diminishes disease-associated excitatory and inhibitory neurons (DAEN and DAIN) in the hippocampus of PS19/E4 mice. a, UMAP plot of all 38 distinct cell clusters in the hippocampi of 9.5-month-old PS19/E4 mice treated with ASO-NTC, ASO-E-4, or ASO-I3-75 for 4 weeks. **b**, Box plot of the proportion of cells from each ASO treatment group in cluster 27. For all box plots, the lower, middle and upper hinges of the box plots correspond to the 25th, 50th and 75th percentiles, respectively. The upper whisker of the box plot extends from the upper hinge to the largest value no further than 1.5 × IQR from the upper hinge. IQR, interquartile range or distance between 25th and 75th percentiles. The lower whisker extends from the lower hinge to the smallest value at most 1.5 × IQR from the lower hinge. Data beyond the end of the whiskers are outlier points. The log odds ratios are the mean ± s.e.m. estimates of log odds ratio for this cluster, which represents the change in the log odds of cells per sample from PS19/E4 mice treated with ASO-E-4 or ASO-I3-75 belonging to the selected cell cluster compared to the log odds of cells per sample from PS19/E4 mice treated with ASO-NTC ASO. **c**, Heat map plot of the log odds ratio per unit change in each pathological parameter for cell clusters 22, 27, and 31. The log odds ratio represents the mean estimate of the change in the log odds of cells per sample from a given treatment group, corresponding to a unit change in a given histopathological parameter. Negative associations are shown in blue and positive associations are shown in red. Unadjusted P values are from fits to a GLMM_histopathology. **d,e,** Volcano plot of the DEGs in excitatory neuron cluster 27 between PS19/E4 mice treated with ASO-E-4 and those treated with ASO-NTC (**d**) and between PS19/E4 mice treated with ASO-I3-75 and those treated with ASO-NTC (**e**). Up- or down-regulated genes shared between **d** and **e** are indicated in red font. **f**, UMAP plot of 23 inhibitory neuron subclusters after subclustering hippocampal cell clusters 8, 11, 15, and 26. **g**,**h**, Box plot of the proportion of cells from each ASO treatment group in inhibitory neuron subcluster 22 (**g**) and inhibitory neuron subcluster 15 (**h**). **i**, Heatmap plot of the log odds ratio per unit change in each pathological parameter for inhibitory neuron subclusters 15 and 22. Negative associations are shown in blue and positive associations are shown in red. Unadjusted P values are from fits to a GLMM_ histopathology. **j,k**, Volcano plot of the DEGs in inhibitory neuron cluster 22 between PS19/E4 mice treated with ASO-E-4 and those treated with ASO-NTC (**j**) and between PS19/E4 mice treated with ASO-I3-75 and those treated with ASO-NTC (**k**). Up- or down-regulated genes shared between **j** and **k** are indicated in red font.

To understand how pathology relates to transcriptomic results, we used log odds ratio (LOR) estimates from a generalized linear mixed-effects model (GLMM) to assess associations with histopathology (GLMM_histopathology). This analysis revealed various excitatory neuron clusters that exhibited significant correlations with disease pathology (Fig. 5c, Extended Data Fig. 4c, Supplementary Table 1). Cell clusters 22, 27, and 31 exhibited significant negative associations with hippocampal volume and positive associations with certain gliosis measurements (Fig. 5c, Extended Data Fig. 4c, Supplementary Table 1). For instance, cluster 22 and 27 were positively correlated with the coverage area of GFAP and APOE4 in neurons (Fig. 5c, Extended Data Fig. 4c), while cluster 31 was positively correlated with coverage area of GFAP, S100β, and CD68 (Fig. 5c, Extended Data Fig. 4c). These associations suggest that these excitatory neuron clusters are disease-associated excitatory neuron (DAEN) subtypes.

To further examine cell type-specific alterations in PS19/E4 mice treated with the different APOE-targeting ASOs, we also used LOR estimates from a GLMM to assess association with animal models (GLMM_AM). Such analysis revealed cell clusters in which there was a significant change in the likelihood that the cluster contained more or fewer cells from PS19/E4 mice treated with ASO-E-4 or ASO-I3-75 than from the mice treated with ASO-NTC. Excitatory neuron clusters 22, 27, and 31 had significantly lower odds of having cells from PS19/E4 mice treated with ASO-E-4 or ASO-I3-75 than from the mice treated with ASO-NTC (Fig. 5b, Extended Data Fig. 4a,b), suggesting diminished DAEN in mice with either global or neuron-preferential reduction of APOE4.

Based on differentially expressed gene (DEG) analyses, cells in neuron clusters 22, 27, and 31 had significantly upregulated expression of the following genes relative to the other excitatory neuron clusters: *Pde4b, St18, Prr5l, Sox2ot, Neat1, Plcl1, Erbin, Plxdc2, Cdk19*, and *Rtn4* (Extended Data Fig. 4d, Extended Data Fig. 5a-c, Supplementary Table 1). Interestingly, *Pde4b* has been implicated in AD in previous studies where inhibition offered protective effects^28^. Notably, DAEN clusters 22, 27, and 31 from PS19/E4 mice treated with either ASO-E-4 or ASO-I3-75 had remarkably similar top upregulated and top downregulated genes when compared to mice treated with ASO-NTC (Fig. 5d,e, Extended Data Fig. 5d-g). This suggests that ASO-E-4 and ASO-I3-75 induce similar transcriptomic changes in these clusters of DAEN, supporting the conclusion that the DEGs induced by global APOE4 reduction in DAEN of PS19/E4 are primarily driven by neuronal APOE4 reduction.

### ASO-E-4 and ASO-I3-75 diminishes disease-associated inhibitory neurons in the hippocampus of PS19/E4 mice

To gain deeper insights into the effects of the two APOE-targeting ASOs on transcriptomics in inhibitory neurons, we did further subclustering analyses of inhibitory neurons in the snRNA-seq dataset.

Subclustering of inhibitory neurons (clusters 8, 11, 15, and 26 in Fig. 5a) identified 23 inhibitory neuron subpopulations (Fig. 5f, Supplementary Table 2). LOR estimates from a GLMM_histopathology revealed that the proportion of cells in inhibitory neuron subclusters 15 and 22 exhibited a significant negative association with hippocampal volume and a significant positive association with neuronal APOE coverage area in the hippocampus (Fig. 5i, Extended Data Fig. 6a, Supplementary Table 2). Additionally, cluster 22 exhibited a significant positive association with markers of gliosis, such as GFAP and CD68 coverage area (Fig. 5i, Extended Data Fig. 6a). Together, these data suggest that inhibitory neuron subclusters 15 and 22 represent disease-associated inhibitory neuron (DAIN) subtypes.

LOR estimates from a GLMM_AM revealed that inhibitory neuron subclusters 15 and 22 had significantly lower odds of having cells from PS19/E4 mice treated with ASO-E-4 or ASO-I3-75 than from the mice treated with ASO-NTC (Fig. 5g,h), suggesting that either global or neuron-preferential reduction of APOE4 diminishes these DAIN subtypes. Based on DEG analyses, cells in inhibitory neuron subclusters 15 and 22 had significantly upregulated expression of the following genes relative to the other inhibitory neuron clusters: *Kcnip4, Pex5l, Marchf1, Sorcs3, Adarb2, Ptprd, Thsd7b, St6galnac3, Brinp3,* and *Nkain2* (Extended Data Fig. 6b-d, Supplementary Table 2). Interestingly, *Kcnip4, Pex5l,* and *Nkain2* are genes important for neuronal excitability; hyperexcitability and network dysfunction are established hallmarks of AD^29–31^. Comparison of DEGs in DAIN subcluster 15 between PS19/E4 mice treated with either ASO-E-4 or ASO-I3-75 and the mice treated with ASO-NTC revealed some similar top downregulated genes, but very few shared top upregulated genes (Extended Data Fig. 6e,f). In a similar DEG comparison for DAIN subcluster 22, PS19/E4 mice treated with either ASO-E-4 or ASO-I3-75 versus mice treated with ASO-NTC had very few shared top downregulated and upregulated genes (Fig. 5j,k). Taken together, these data illustrate that both global and neuron-preferential reduction of APOE4 by APOE-targeting ASOs diminishes DAIN. However, the transcriptomic changes induced by the global APOE4 reduction and the neuron-preferential APOE4 reduction are different in these DAINs, particularly DAIN subcluster 22, likely contributing to their differential effects on rescuing hippocampal neurodegeneration.

### ASO-I3-75 is more effective than ASO-E-4 at reducing disease-associated astrocytes in the hippocampus of PS19/E4 mice

We next wanted to better understand the effects of the two APOE-targeting ASOs on astrocytes. To do so, we further subclustered the astrocytes of the snRNA-seq dataset (cluster 13 in Fig. 5a) and identified 15 astrocyte subpopulations (Fig. 6a, Supplementary Table 3). LOR estimates from a GLMM_histopathology revealed that the proportion of cells in astrocyte subclusters 4 and 9 exhibited a significant negative association with hippocampal volume and a significant positive association with GFAP coverage area (Fig. 6d, Extended Data Fig. 7a, Supplementary Table 3). Subcluster 4 also had a significant negative association with DG neuronal layer thickness and a significant positive association with S100β and CD68 coverage area; and subcluster 9 exhibited a positive association with IBA1 and hippocampal neuronal APOE coverage area (Fig. 6d, Extended Data Fig. 7a, Supplementary Table 3). Altogether, these associations suggest that astrocyte subclusters 4 and 9 represent disease-associated astrocyte (DAA) subtypes.

**Fig. 6.**
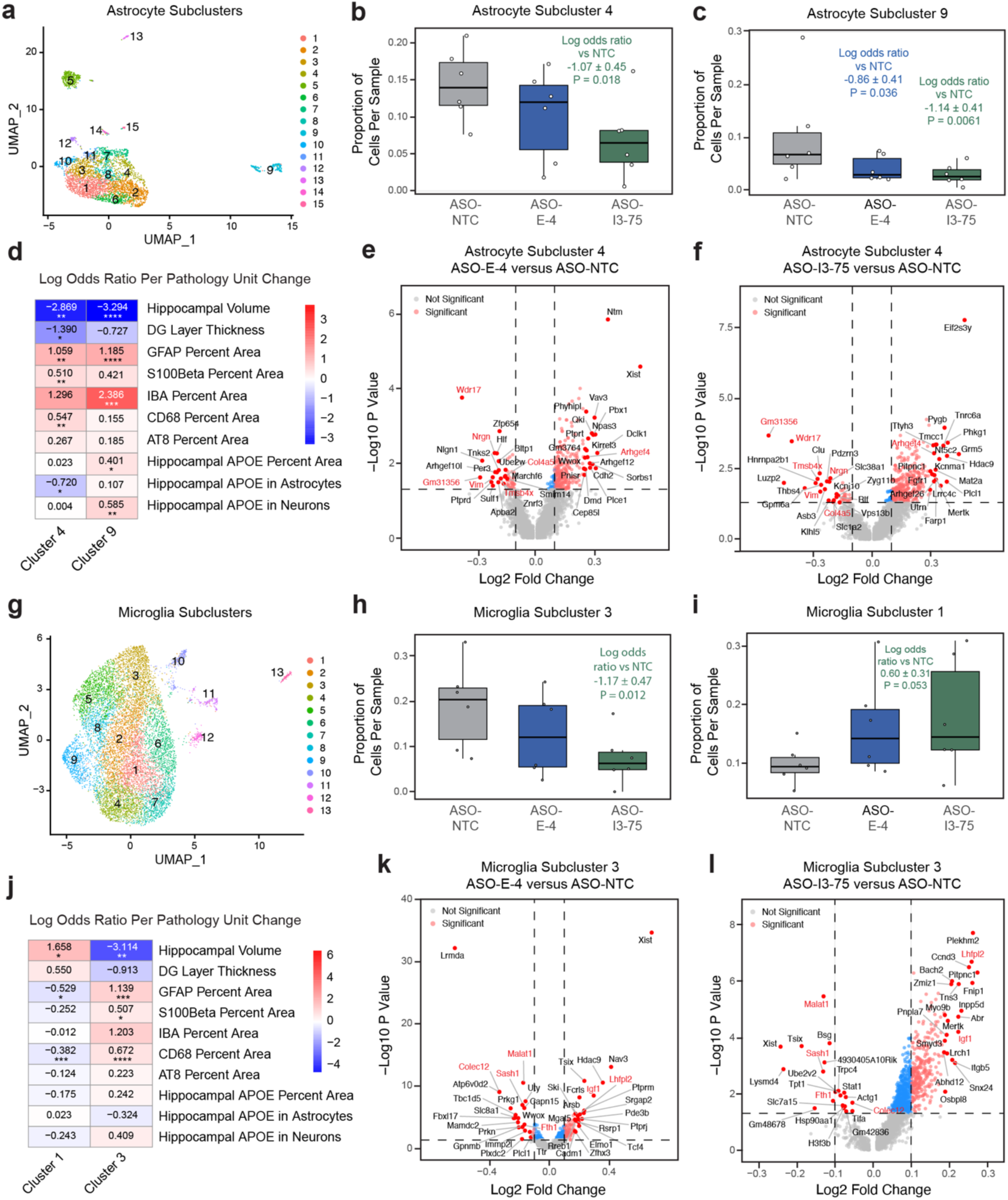
ASO-I3-75 is more effective than ASO-E-4 at reducing disease-associated astrocytes (DAA), reducing disease-associated microglia (DAM), and increasing disease-protective microglia (DPM) in the hippocampus of PS19/E4 mice. **a**, UMAP plot of 15 astrocyte subclusters after subclustering hippocampal cell cluster 13. **b,c**, Box plot of the proportion of cells from each ASO treatment group in astrocyte subcluster 4 (**b**) and astrocyte subcluster 9 (**c**). For details see legend in Fig. 5b. **d**, Heatmap plot of the log odds ratio per unit change in each pathological parameter for astrocyte subclusters 4 and 9. Negative associations are shown in blue and positive associations are shown in red. Unadjusted P values are from fits to a GLMM_ histopathology. **e,f**, Volcano plot of the DEGs in astrocyte subcluster 4 between PS19/E4 mice treated with ASO-E-4 and those treated with ASO-NTC (**e**) and between PS19/E4 mice treated with ASO-I3-75 and those treated with ASO-NTC (**f**). Up- or down-regulated genes shared between **e** and **f** are indicated in red font. **g**, UMAP plot of 13 microglia subclusters after subclustering hippocampal cell clusters 6 and 21. **h,i**, Box plot of the proportion of cells from each ASO treatment group in microglia subclusters 3 (**h**) and 1 (**i**). For details see legend in Fig. 5b. **j**, Heatmap plot of the log odds ratio per unit change in each pathological parameter for microglia subclusters 1 and 3. Negative associations are shown in blue and positive associations are shown in red. Unadjusted P values are from fits to a GLMM_ histopathology. **k,l**, Volcano plot of the DEGs in microglia subcluster 3 between PS19/E4 mice treated with ASO-E-4 and those treated with ASO-NTC (**k**) and between PS19/E4 mice treated with ASO-I3-75 and those treated with ASO-NTC (**l)**. Up- or down-regulated genes shared between **k** and **l** are indicated in red font.

LOR estimates from a GLMM_AM revealed that astrocyte subcluster 4 had significantly lower odds of having cells from PS19/E4 mice treated with ASO-I3-75 than from the mice treated with ASO-NTC (Fig. 6b), suggesting that neuron-preferential reduction of APOE4 is particularly effective at diminishing this DAA subtype. Interestingly, astrocyte subcluster 9 had significantly lower odds of having cells from PS19/E4 mice treated with either ASO-E-4 or ASO-I3-75 than from the mice treated with ASO-NTC (Fig. 6c), suggesting that either global or neuron-preferential reduction of APOE4 is effective at diminishing this particular DAA subtype.

DEG analyses revealed that cells in DAA subclusters 4 and 9 had significantly upregulated expression of the following genes relative to the other astrocyte clusters: *Grm5, Ncam2, Enox1, Chl1, Lrp1b, Cadm2, Nav2, Rfx3, Stxbp6,* and *Cadps* (Extended Data Fig. 7b, Extended Data Fig. 8a,b, Supplementary Table 3). Comparison of DEGs in DAA subcluster 4 between PS19/E4 mice treated with ASO-E-4 or ASO-I3-75 and the mice treated with ASO-NTC revealed some similar top downregulated genes, but very few shared top upregulated genes (Fig. 6e,f). This suggests that the upregulation of genes induced by the global APOE4 reduction and the neuron-preferential APOE reduction are different in DAA subcluster 4, likely contributing to their differential effects on rescuing hippocampal neurodegeneration. Interestingly, DEG comparison in subcluster 9 between PS19/E4 mice treated with ASO-E-4 or ASO-I3-75 and the mice treated with ASO-NTC yielded many similar top upregulated and downregulated genes (Extended Data Fig. 8e,f). This suggests that ASO-E-4 and ASO-I3-75 induce similar transcriptomic changes in DAA subcluster 9, supporting the conclusion that the DEGs induced by global APOE4 reduction in this subtype of DAA in PS19/E4 mice are primarily driven by neuronal APOE4 reduction. Taken together, all these data illustrate that both global and neuron-preferential reduction of APOE4 diminishes subtypes of DAA in PS19/E4 mice. However, neuron-preferential reduction of APOE4 seems to be more effective at reducing multiple subtypes of DAA, probably contributing to its more effective rescue of hippocampal neurodegeneration in this tauopathy mouse model.

### ASO-I3-75, but not ASO-E-4, reduces disease-associated microglia and increases disease-protective microglia in the hippocampus of PS19/E4 mice

Lastly, to better understand the effects of the two APOE-targeting ASOs on microglia, we further subclustered microglia (cluster 6 and 21 in Fig. 5a), identifying 13 microglia subpopulations (Fig. 6g, Supplementary Table 4). LOR estimates from a GLMM_histopathology revealed that the proportion of cells in microglia subcluster 3 exhibited a significant negative association with hippocampal volume and a significant positive association with GFAP coverage area, S100β coverage area, and CD68 coverage area (Fig. 6j, Extended Data Fig. 7c, Supplementary Table 4), suggesting that microglia subcluster 3 represents a disease-associated microglia (DAM) subtype.

LOR estimates from a GLMM_AM revealed that microglia subcluster 3 had significantly lower odds of having cells from PS19/E4 mice treated with ASO-I3-75 than from the mice treated with ASO-NTC (Fig. 6h), suggesting that neuron-preferential, but not global, reduction of APOE4 diminishes this subtype of DAM. DEG analyses revealed that cells in microglia subcluster 3 had significantly upregulated expression of the following genes relative to the other microglia clusters: *Apobec1, Gpnmb, Lyst, Fnip2, Igf1, Myo5a, Atp6v0d2, Pid1, Colec12,* and *Arhgap15* (Extended Data Fig. 7d, Extended Data Fig. 8c, Supplementary Table 4). *Gpnmb* is an important DAM gene and is known to increase in an age-dependent manner in transgenic AD models^32^. Both *Lyst* and *Atp6v0d2* are important genes for lysosomal function, and lysosomal dysfunction leading to profuse accumulations of neuronal waste is a known pathology in AD^33^. Notably, comparison of DEGs in DAM subcluster 3 between PS19/E4 mice treated with either ASO-E-4 or ASO-I3-75 and the mice treated with ASO-NTC yielded several similar top downregulated genes and very few similar top upregulated genes (Fig. 6k,l). This suggests that the upregulation of genes induced by the global APOE4 reduction and the neuron-preferential APOE reduction are different in DAM subcluster 3, likely contributing to their differential effects on rescuing hippocampal neurodegeneration. Overall, neuron-preferential reduction of APOE4 seems to be more effective at reducing DAM in PS19/E4 mice compared to global reduction of APOE4.

Interestingly, LOR estimates from a GLMM_histopathology also revealed that the proportion of cells in microglia subcluster 1 exhibited a significant positive association with hippocampal volume and a significant negative association with GFAP coverage area and CD68 coverage area (Fig. 6j, Extended Data Fig. 7c, Supplementary Table 4), suggesting that microglia subcluster 1 represents a disease protective microglia (DPM) subtype.

LOR estimates from a GLMM_AM revealed that DPM subcluster 1 had strong trending higher odds (p = 0.053) of having cells from PS19/E4 mice treated with ASO-I3-75 than from the mice treated with ASO-NTC (Fig. 6i). Based on DEG analyses, cells in microglia subcluster 1 had significantly upregulated expression of the following genes relative to the other microglia clusters: *Malat1, Ttr, Tmsb4x, Arhgap45, Gpr34, Dynll1, Tanc2, Siglech, St3gal5,* and *Rhobtb1* (Extended Data Fig. 7d, Extended Data Fig. 8d, Supplementary Table 4). Interestingly, DPM subcluster 1 had significant downregulated expression of the *Apobec1*, *Myo5a*, *Fnip2*, *Lyst* and *Arhgap15 –* genes that were upregulated in DAM subcluster 3—further supporting the conclusion that it is a DPM subtype (Extended Data Fig. 7d, Extended Data Fig. 8c,d). Furthermore, *Tanc2*, which is upregulated in DPM subcluster 1, is downregulated in DAM subcluster 3 (Extended Data Fig. 7d, Extended Data Fig. 8c,d). Comparing DEGs in DPM subcluster 1 between PS19/E4 mice treated with either ASO-E-4 or ASO-I3-75 and the mice treated with ASO-NTC yielded similar top upregulated and top downregulated DEGs (Extended Data Fig. 8g,h, Supplementary Table 4). This suggests that ASO-E-4 and ASO-I3-75 induce similar transcriptomic changes in this DPM subtype, supporting the conclusion that the DEGs induced by global APOE4 reduction in this DPM subtype in PS19/E4 mice are primarily driven by neuronal APOE4 reduction. However, neuron-preferential reduction of APOE4 appears to be more effective than global reduction of APOE4 at increasing this DPM subtype. Taken together, these data illustrate that neuron-preferential reduction of APOE4 is more effective than global reduction of APOE4 at decreasing DAM and increasing DPM in PS19/E4 mice.

## DISCUSSION

In this study, we investigated the therapeutic benefit of decreasing APOE4 expression globally or preferentially in neurons in the brain of an AD mouse model via the use of ASOs. We designed an ASO targeting APOE produced by all cell types in the brain (ASO-E-4) and an ASO preferentially targeting APOE produced by neurons (ASO-I3-75). First, we validated the efficacy of these ASOs *in vitro* using various cell culture systems and determined that ASO-I3-75 does preferentially reduce neuronal APOE by targeting a neuron-specific APOE-I3 transcript. Then, we demonstrate that treating PS19/E4 mice with ASO-I3-75 significantly rescues hippocampal neurodegeneration, alters p-tau contribution to neurodegeneration, decreases astrogliosis, alters microgliosis contribution to neurodegeneration, diminishes disease-associated excitatory and inhibitory neurons, reduces disease-associated astrocytes and microglia, and increases disease-protective microglia. While treating PS19/E4 mice with the global brain APOE-targeting ASO, ASO-E-4, recapitulates some of the abovementioned beneficial effects, such as decreasing astrogliosis and diminishing disease-associated neurons, it does not recapitulate all of them. Overall, these findings illustrate that preferentially reducing neuronal APOE4 is more beneficial than globally reducing APOE4 in the brain for protecting against or rescuing key AD-related pathologies.

*In vitro* studies using mouse neuroblastoma cells stably transfected with human APOE3 genomic DNA and mouse primary neurons expressing human APOE4 from a mini-APOE4 gene (containing all exons and intron-3) demonstrated that treatment with either ASO-E-4 or ASO-I3-75 effectively reduces neuronal APOE expression. This is likely because the exon-4-targeting ASO-E-4 targets both mature APOE transcripts without intron-3 and neuron-specific APOE-I3 transcripts, whereas the intron-3-targeting ASO-I3-75 only targets the neuron-specific APOE-I3 transcripts, but not mature APOE transcripts. Similarly, *in vivo* studies with PS19/E4 mice show that both ASOs effectively reduce APOE4 expression in neurons. Therefore, the decreased astrogliosis and diminished disease-associated excitatory neurons that we observed following treatment with the global brain APOE-targeting ASO, ASO-E-4, could likely be achieved, at least in part, by decreasing neuronal APOE4 expression.

PS19/E4 mice treated with the neuronal-APOE-preferential ASO, ASO-I3-75, exhibited a significantly larger hippocampal volume compared to PS19/E4 mice treated with ASO-NTC, suggesting that preferentially reducing neuronal APOE4 effectively rescues hippocampal neurodegeneration. Strikingly, a significant negative correlation was observed between neuronal APOE4 levels and hippocampal volume in mice treated with ASO-I3-75, but not in mice treated with ASO-E-4 or ASO-NTC. This suggests that in a context where we can successfully decrease neuronal APOE4 expression levels, the extent of APOE4 reduction in neurons is a good predictor for the efficacy of hippocampal volume rescue.

When correlating hippocampal volume to gliosis measurements, such as S100β percent coverage area, IBA1 percent coverage area, and CD68 percent coverage area, a significant or trending negative correlation was observed in mice treated with ASO-NTC or ASO-E-4, suggesting that, in these mice, the severity of gliosis is a good predictor for hippocampal volume loss. Interestingly, for all three gliosis measurements in mice treated with ASO-I3-75, there was no longer a significant negative correlation with hippocampal volume, suggesting that in mice with neuronal APOE4 being preferentially reduced, higher levels of gliosis are no longer a good predictor for hippocampal volume loss. One possible explanation is that ASO-I3-75 treatment increases disease-protective microglial or astrocytic subpopulations, contributing to the increase in hippocampal volume. An example of such a disease-protective microglial subtype is the DPM subcluster 1.

A limitation of our current study is that we were unable to directly compare neuronal-APOE ASO reduction with an astrocyte-specific or microglia-specific APOE ASO reduction approach, largely due to the lack of technology enabling glial-specific ASO delivery. However, reducing glial APOE could confer protection against some AD-related pathologies. Studies have found that removal of APOE4 from astrocytes in a tauopathy mouse model reduces the extent of tau pathology, neurodegeneration, and gliosis^27^. To get a better understanding of this, further therapeutic studies should investigate cell type-specific APOE4 reduction in astrocytes or microglia starting at a similar age and in a similar PS19/E4 mouse model to see if reduction of APOE4 in only these cell types rescues AD pathologies. Since traditional ASO designs are not feasible to specifically target astrocytes or microglial APOE4, unique approaches are needed to reach this goal in the future.

It is also worth noting that PS19/E4 mice treated with ASO-I3-75 exhibited a noticeable decrease, despite not reaching statistical significance, in astrocytic APOE as well. This could be due to an effect of the ASO-I3-75 binding to some pre-mRNAs in astrocytes that contain intron-3 and thus, decreasing their conversion into mature APOE mRNA. Alternatively, given that previous studies found intron-3 containing APOE transcripts are not abundant in non-neuronal cell types^20^, this reduction could instead be due to secondary effects of the ASO-I3-75 efficiently decreasing neuronal APOE4. Decreased neuronal APOE4 could result in a decrease in astrocyte response (astrogliosis) to neuronal damage, and thus a decrease in astrocytic APOE4 because the numbers of astrocytes have decreased or their APOE4 expression levels have been reduced.

We reported previously that selective neuronal APOE4 removal in PS19/E4 mice from birth led to a wide range of beneficial effects, including reductions in the accumulation and spread of p-tau, neurodegeneration, myelin deficits, neuronal network hyperexcitability, microgliosis, astrogliosis, and disease-associated cell subpopulations^11^. In the present study, we expand on this body of work and demonstrate that preferentially reducing neuronal APOE4 by ASO-I3-75 treatment in 8.5-month-old PS19/E4 mice, after pathologies have initiated, also protects against and recues APOE4-driven AD pathologies.

Furthermore, studies have found that global depletion of APOE4 may have deleterious side effects^34–38^, suggesting that it would be more beneficial to achieve specific reduction of APOE4 in a certain cell type most vulnerable to APOE4 detrimental effects, such as neurons, while leaving the physiological functions of APOE4 largely intact in other types of cells. Our current study provides a proof-of-concept case showing that preferentially reducing neuronal APOE4 with ASO-I3-75 treatment can eliminate most of the pathogenic effects of APOE4 while largely leaving APOE4 intact in other cells for its physiological functions. Clearly, further preclinical studies need to be done to better understand the underlying mechanisms and to identify optimal treatment dose and duration for potential clinical trial tests in AD patients with APOE4 in the future.

## METHODS

### Mice

Human LoxP-floxed APOE4 knock-in (fE4) mice^39^ and tau-P301S fE4 (PS19-fE4) mice^11,25^ were generated as previously described. All mice were on a pure C57BL/6 genetic background and were housed in a pathogen-free barrier facility on a 12-h light cycle at 19–23°C and 30–70% humidity. Animals were identified by ear punch and genotyped by TransnetYX automated genotyping PCR services using tail clippings. For all studies, both male and female mice were used. All animal experiments were conducted in accordance with the guidelines and regulation of the National Institutes of Health, the University of California and the Gladstone Institutes under the protocols AN176773 and AN195565. All protocols and procedures followed the guidelines of the Laboratory Animal Resource Center at the University of California, San Francisco (UCSF) and the ethical approval of the UCSF institutional animal care and use committee.

Mice were sedated with avertin and transcardially perfused with sterile saline. The right hemi-brains were drop-fixed for 48 h in 4% paraformaldehyde (Electron Miscropscopy Science, Cat. #15710-S), washed for 24 h in 1X PBS (Thermo Scientific, Cat. #J75889K2) and cryoprotected in 30% sucrose (Millipore Sigma, Cat. #S7903) for 48 h at 4°C. The fixed right hemispheres were cut into 30-μm thick coronal sections on a freeze sliding microtome (Leica, Cat. #SM2010R) and stored in cryoprotectant solution at −20°C (30% ethylene glycol, Fisher Chemical, Cat. #E178-4; 30% glycerol, Millipore Sigma, Cat. #G9012-1GA; and 40% 1x PBS). Left hemispheres were snap-frozen on dry ice and stored at−80°C.

### Antisense oligonucleotides (ASOs)

The ASOs used were synthesized by Integrated DNA Technologies (IDT). They were designed to have a central core of phosphorothioate-modified DNA flanked by 2′-O-methoxyethyl (2′-MOE) modified RNA bases. Specific modifications include: 5 nucleotides on the 5′ and 3′ termini containing 2′-MOE modifications, 10 unmodified central oligodeoxynucleotides, and a phosphorothioate backbone. These modifications were incorporated to enhance nuclease resistance, lower cell toxicity, increase binding affinity to the desired target, and promote cellular uptake. For *in vitro* and *ex vivo* studies, ASOs were solubilized in 10 mM Tris, 0.1 mM EDTA (TE) buffer (Thermo Scientific Chemicals, Cat. #J75793.AP). For *in vivo* studies, ASOs were HPLC-purified, underwent sodium-salt exchange, and were solubilized in 0.9% sterile saline (Hospira, Cat. #0409-4888-02). ASO-targeting regions were as follows:

| ASO Name | Targeting Region in the APOE Gene |
| --- | --- |
| APOE<sup>ASO-4</sup> | Exon-4 |
| APOE<sup>ASO-75</sup> | Intron-3 |

| Non-targeting negative control (NTC) | Non-Targeting Control |
| --- | --- |
| APOE<sup>ASO-133</sup> | Exon-3 |
| APOE<sup>ASO-174</sup> | Exon-3 |
| APOE<sup>ASO-269</sup> | Exon-4 |
| APOE<sup>ASO-842</sup> | Exon-4 |
| APOE<sup>ASO-485</sup> | Intron-3 |
| APOE<sup>ASO-201</sup> | Intron-3 |
| APOE<sup>ASO-431</sup> | Intron-3 |
| APOE<sup>ASO-415</sup> | Intron-3 |

### Stereotaxic surgery and surgical placement of ICV pumps in mice

Mice were anesthetized with isoflurane and maintained on 1.8-2.0% isoflurane (Dechra, Cat. #200-129) for the surgery. After anesthesia, the mice were secured in a stereotaxic frame with a nose cone (Kopf instruments Model 940). The hair on the head was removed and a subcutaneous injection of anesthetic (Lidocaine; Vedco) was injected above the skull where the cut in the scalp was going to be made. After cutting open the scalp with sterilized scissors, cranial sutures were visualized using 3% hydrogen peroxide. Next, a unilateral stereotaxic site was drilled with a 0.5-mm microburr (Fine Science Tools, Cat. #19007-05) to target the right lateral ventricle with the following coordinates based on bregma:−0.25 mm posterior, 1 mm lateral (right), and −2.0 mm ventral. A 28-day Alzet osmotic pump (Model 2004, Cat. #0000298) with ASO was then implanted in a subcutaneous pocket formed on the back of the mice with the catheter placed in the drill site. Following surgery, mice were sutured with nylon monofilament non-absorbable 6-0 sutures (Surgical Specialties Look, Cat. #911B) and administered analgesics buprenorphine (Henry Schein, Cat. #55175) and ketofen (Henry Schein, Cat. #005487). Mice were monitored on a heating pad until ambulatory, provided diet gel, and housed individually with extra enrichment provided.

### Quantitative real-time polymerase chain reaction

RNA analyses were performed using quantitative real-time polymerase chain reaction (qRT-PCR). Total RNA was extracted from cells using a Qiagen RNeasy Micro Kit (Qiagen, Cat. #74004). RNA was reverse-transcribed and cDNA generated using the SuperScript IV First-Strand Synthesis System (Invitrogen, Cat. # 18091050). The qRT-PCRs were run and analyzed using the QuantStudio 5 Real-Time PCR System (Applied Biosystems, Cat. # A28140). Total human APOE and APOE-I3 mRNA expression levels were normalized to mouse glyceraldehyde-3-phosphate dehydrogenase (GAPDH) mRNA levels analyzed using the DDCt method for relative expression analysis. Primers were synthesized by ElimBio and the sequences were as follows:

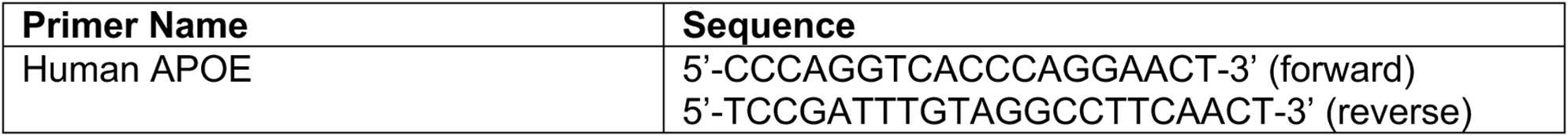
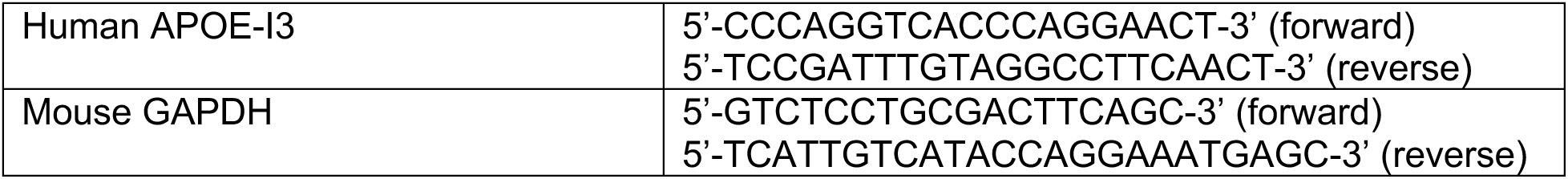

### Cell cultures and ASO treatment

Mouse neuroblastoma Neuro-2a cells stably transfected with human apoE3 genomic DNA that were established as previously described^22^ were used for the initial screen of ASOs. Neuro-2a cells were maintained at 37°C in a humidified 5% CO2 incubator in DMEM, high glucose (Gibco, Cat. #11965092) supplemented with 10% FBS (Gibco, Cat. #A56697-01), 1X nonessential amino acids (NEAA) (Gibco, Cat. #11140050), and 0.5% penicillin/streptomycin (Gibco, Cat. #15140122). For ASO treatments, Neuro-2a were transfected using Lipofectamine 2000 Transfection Reagent (Invitrogen, Cat. #11668019). The day before transfection with ASOs, cells were lifted using 0.05% Trypsin-EDTA (Gibco, Cat. #25300054), counted, and plated at a concentration of 1.5 x 10^5^ cells per well in a 24-well plate using media without antibiotics. For transfection, an Opti-MEM Reduced Serum Media (Gibco, Cat. #31985070) combined with ASO mixture and an Opti-MEM Reduced Serum Media combined with Lipofectamine 2000 mixture were made and incubated at room temperature (RT) for 5 min. The Opti-MEM + ASO mixture was made using various ASOs and various concentrations depending on experiment. The diluted ASO mixture and the diluted Lipofectamine 2000 mixture were next mixed together gently by pipetting up and down and incubated at RT for 25 min. After feeding the Neuro-2a cells with fresh media without antibiotics, the transfection mix was added dropwise into the well and gently rocked back and forth. After ASO treatment, the cells were collected for either qRT-PCR or western blot analysis. For qRT-PCR, cells were lysed directly in the well using Buffer RLT provided in the Qiagen RNeasy Kit supplemented with 2M dithiothreitol (DTT) (Bioworld, Cat. #20840007) at a concentration of 20 uL per mL of RLT. After collection, the samples were snap frozen on dry ice and stored −80 °C. For western blot analysis, cells were lysed in the well using a high-detergent buffer: 50 mM Tris, 150 mM NaCl, 2% NP-40 (Millipore Sigma, Cat. #492016), 1% sodium deoxycholate and 4% sodium dodecyl sulfate and supplemented with complete protease inhibitor cocktail pellet (Roche, Cat. 11697498001), phosphatase inhibitor cocktail 2 (Millipore Sigma, Cat. #P5726), phosphatase inhibitory cocktail 3 (Millipore Sigma, Cat. #P0044) and Benzonase Nuclease (Millipore Sigma, Cat. #E1014-25KU). After collection, the samples were centrifuged at maximum speed for 5 min and the supernatant was collected and stored at −80°C.

Human astrocytoma cells, SW-1088, were maintained at 37°C in a humidified 0% CO2 incubator in Leibovitz’s L-15 Medium (Gibco, Cat. #11415064) supplemented with 10% FBS and 1% penicillin/streptomycin. ASOs were introduced to cells via gymnotic delivery. The day before treatment with ASOs, cells were lifted using 0.05% Trypsin-EDTA and plated at a concentration of 1.5 x 10^5^ cells per well in a 24-well plate. ASOs were diluted from 100 μM stock concentration to 3 μM with 500 uL of media per well. After 48 h, SW-1088 cells were collected for qRT-PCR via the same method as described above for Neuro-2a cells.

### Primary neuron culture and ASO treatment

Primary neuron culture was prepared from post-natal day 0 (P0) pups as previously described^12^. Briefly, after removing the head from the pup, the brain was dissected out and placed into a dish with ice-cold dissociation media (DM) containing kynurenic acid (KY), called DM/KY media. More specifically, DM/KY is made up of 90% DM and 10% KY stock. DM: 81.8 mM Na_2_SO_4_ (Millipore Sigma, Cat. #S5640), 30 mM K_2_SO_4_ (Millipore Sigma, Cat. #P8541), 5.8 mM MgCl_2_ (Millipore Sigma, Cat. #M1028), 0.25 mM CaCl_2_ (Millipore Sigma, Cat. #C5670), 1 mM HEPES (Millipore Sigma, Cat. #H0887), 20 mM Glucose (Millipore Sigma, Cat. #G8769), 0.001% Phenol Red (Millipore Sigma, Cat. #P-0290), 0.16 mM NaOH (Ricca Chemical Cat. #7470-32). KY: 10 mM kynurenic acid (Millipore Sigma, Cat. #K3375), 0.0025% Phenol Red, 5 mM HEPES, 100 mM MgCl_2_, NaOH to titrate pH to 7.4. With the dissected brain, the meninges were removed and the cortex was isolated using a dissection microscope. Next, the tissue was minced using a sterilized razor blade and digested using a Papain (Worthington, Cat. #LS003124) solution. The Papain solution was made by diluting Papain in DM/KY such that there were 5.5 units per mL of DM/KY. After digestion at RT for 13 min, with gentle tube inversions every couple of minutes, the Papain solution was removed, and a Trypsin inhibitor solution was added. The Trypsin inhibitor solution was made by diluting 1 g of Trypsin inhibitor (Millipore Sigma, Cat. #T9253) with 100 mL of DM/KY and titrating the pH to 7.4 with NaOH. After two washes with the Trypsin inhibitor solution for 5 min, the tissue pellet was washed twice with an Opti-MEM/Glucose solution for 5 min. During the second wash with the Opti-MEM/Glucose solution, the tissue pellet was gently triturated using a 5 mL pipette into a single cell suspension. The Opti-MEM/Glucose solution was made by adding Glucose to Opti-MEM (Thermo) to be a total concentration of 20 mM. The cells were then spun down at 200G for 5 min, resuspended in B27/Neurobasal growth media, counted, and plated at a concentration of 6.0 x 10^5^ cells per well in a 24-well plate. The B27/Neurobasal growth media consisted of Neurobasal medium (Gibco, Cat. #21103-049), 1X B27 (Gibco, Cat. #17504-044), 1% Glutamax (Gibco, Cat. #35050061), and 1% penicillin/streptomycin (Gibco, Cat. #15140122). Prior to plating cells on the 24-well plates, the wells were coated with PLL (Millipore Sigma, Cat. #P4707) and washed three times with DPBS (Thermo Scientific Chemicals, Cat. #J67653AP). ASOs were introduced to primary neurons via gymnotic delivery. The primary neuron cultures were treated with 10 μM of ASO on Day 3 for 1 week and collected on Day 10 for qRT-PCR via the same method as described above for Neuro-2a cells.

### Primary astrocyte culture and ASO treatment

Mouse primary mixed glia culture was prepared from neonatal mice at P2-3 from PS19/E4 mice as previously described^40^. Briefly, the cortices were separated from the meninges and minced with a sterile razor blade. The tissue was digested with 2.5% trypsin (Gibco, Cat. # 15090046) + DNase (Millipore Sigma, Cat. #D4513) for 25 minutes at 37°C. After incubation, the trypsin was neutralized with 30% FBS (Gibco, Cat. #A56697-01) in DMEM, high glucose, GlutaMAX Supplement, pyruvate (Gibco, Cat. #10569010) and triturated until cells formed a uniform suspension. The cells were spun at 300G for 10 min and pellet was resuspended in 10% FBS in DMEM. For each brain from a single pup, cells were plated onto PDL (Millipore Sigma, Cat. #P7280) coated T25 flasks supplemented with 50ng/mL GM-CSF (Millipore Sigma, Cat. # G0282). On Day 4, the fresh media was added onto the cells. On Day 10-12, the flasks containing mixed glial were placed onto an incubated shaker at 37°C for 1 h at 180RPM to shake off the microglia. After the microglia were shaken off, the remaining astrocytes in the T25 flasks were fed with fresh media and kept at 37°C until they were lifted and replated onto 24-well plates. To replate astrocytes, the media was removed and 3 mL of 2.5% trypsin was added to the flask. Next the cells were incubated at 37°C for 5 min until the cells lifted up from the surface of the flask. Then, using a 5 mL pipette, the cells were pipetted up and down a few times to break down the sheet of cells before being incubated at 37°C for another 5 min. After this, the cells were triturated using a 5 mL pipette until the cells were a uniform, cloudy solution. An equal volume of growth media was then added to the cells to quench the trypsin before the cell solution was passed through a 40 mm filter (Corning, Cat. #431750). The cells were then spun at 300G for 5 min, the cell pellet resuspended in media, and plated at a concentration of 3.0 x 10^5^ cells per well in a 24-well plate. Prior to plating primary astrocytes on 24-well plates, the wells were coated with PLL (Millipore Sigma, Cat. #P4707) and washed three times with DPBS (Thermo Scientific Chemicals, Cat. #J67653AP). ASOs were introduced to primary astrocytes via gymnotic delivery. The primary astrocytic cultures were treated with 20 or 40 μM of ASO on Day 14 for 1 week and collected on Day 21 for qRT-PCR via the same method as described above for Neuro-2a cells.

### Immunohistochemistry

For immunofluorescence staining, 30 μm-thick brain sections were washed three times with PBS-T (PBS + 0.1% Tween-20; Millipore Sigma, Cat. #P2287) and incubated in boiling 1M Tris-HCl buffer (pH 8, Corning, Cat. #46-031-CM) for 5 min. Next, sections were washed three times with PBS-T followed by blocking with 10% normal donkey serum (NDS; Jackson ImmunoResearch, Cat. #017000121) made in PBS-Tx (PBS + 0.5% TritonX; Millipore Sigma, Cat. #T8787) for 1 hr. Sections were then incubated in Mouse on Mouse (MOM; Vector Laboratories, Cat. #MKB-2213-1) blocking buffer (one drop MOM per 4ml of PBS) to block endogenous mouse staining. Then, the sections were incubated overnight at 4°C with primary antibodies. The following primary antibodies were used for studies: anti-APOE (rb) 1:500 (Cell Signaling Technology, Cat. #13366); anti-NeuN 1:500 (Millipore Sigma, Cat. #ABN90); anti-GFAP 1:750 (Millipore Sigma, Cat. #MAB3402); anti-S100B 1:200 (Abcam, Cat. #ab52642); anti-IBA1 (gt) 1:500 (Abcam, Cat. #ab5076); and anti-CD68 1:200 (Bio-Rad, Cat. #MCA1957). All primary antibodies were diluted in 3% NDS made in PBS. The next day, sections were washed three times with PBS-T and incubated in the dark for 1 hour at room temperature with secondary antibodies. The following secondary antibodies were used: donkey anti-rabbit 488 1:1000 (Invitrogen, Cat. #A-21206); donkey anti-guinea pig 647 1:1000 (Jackson Immuno, Cat. #706-605-148); donkey anti-mouse 594 1:1000 (Invitrogen, Cat. #A-21203); donkey anti-rabbit 647 1:1000 (Invitrogen, Cat. #A231573); donkey anti-goat 594 1:1000 (Invitrogen, Cat. #A11058); donkey anti-rat 594 (Invitrogen, Cat. #A21209). All secondary antibodies were diluted in 3% NDS made in PBS. During the incubation with secondary antibodies, for some stains, DAPI (1:20,000; Thermo Scientific, Cat. #62248) was added. After incubation with secondary antibodies, the sections were then washed three times with PBS-T before getting mounted onto microscope slides (Fisher Scientific, Cat. #22-037-246). Slides were coverslipped with ProLong Gold mounting medium (Invitrogen, Cat. #P36930), allowed to dry, and followed by imaging on a FV3000 confocal laser scanning microscope (Olympus).

For diaminobenzidine (DAB) staining, 30um-thick brain sections were washed three times with PBS-T, incubated in boiling citrate buffer (10mM sodium citrate; Fisher Bioreagents, Cat. # BP327-1) for 5 min, washed two times with PBS-T, incubated in endogenous peroxidase block (3% H2O2; Millipore Sigma, Cat. #H1009-500mL, and 10% methanol; Fisher Scientific, Cat. #A412SK-4, in PBS) for 15 min, washed two more times with PBS-T, washed one time with PBS-Tx (PBS + 0.2% TritonX; Millipore Sigma, Cat. #T8787), and blocked with 10% NDS made in PBS-Tx for 1 hr. Sections were then incubated in avidin and biotin blocks (3 drops of each; Vector Laboratories, Cat. #SP-2001) for 15 min and MOM Blocking buffer (one drop MOM per 4 ml of PBS-T) for 1 hr. Sections were incubated overnight at 4°C with an anti-AT8 primary antibody 1:100 (Invitrogen, Cat. #MN1020) diluted in 3% NDS made in PBS-T. The following day, sections were washed three times with PBS-T and incubated for 1 hr at room temperature with biotinylated donkey anti-mouse secondary (1:2000; Jackon ImmunoResearch, Cat. #715-065-150) diluted in 3% NDS in PBS-T. Next, the sections were rinsed two times with PBS-T, once with PBS, and incubated with ABC buffer (2 drops A and B per 5ml PBS; Vector Laboratories, Cat. #PK-6100) for 1 hr. The ABC buffer was made and allowed to incubate 15-30 min before use. Next, the sections were rinsed twice with PBS and once with water before a 1 min and 30 sec incubation in DAB buffer (2 drops buffer stock solution, 4 drops DAB, and 2 drops H_2_O_2_ in 5ml milliQ water; Vector Laboratories, Cat. #SK-4100). Sections were then immediately washed three times with milliQ water and then washed with PBS and mounted onto microscope slides. Slides were allowed to dry overnight and then submerged in two washes of xylene (Fisher Scientific, Cat. #HC7001GAL). Slides were coverslipped with DPX mounting medium (Millipore Sigma, Cat. #06522) and imaged with an Aperio VERSA slide scanning microscope (Leica).

### Immunohistochemical quantification analysis

For all image quantifications, researchers were blinded to samples and automated imaging analysis via Fiji was used to exclude bias from quantifications. For analyses where immunostaining as a percentage of hippocampal area covered was conducted, two brain sections for each mouse was stained and imaged according to the immunohistochemistry section above. Next, four fields of view from the two brain sections (two fields of view from the DG and two fields of view from the CA1) were taken and used to calculate an average percent area coverage for each mouse.

For percent APOE area coverage in astrocytes and neuron quantifications, brain sections were stained according to the immunohistochemistry section above with anti-GFAP or anti-NeuN to highlight astrocytes or neurons, respectively. Next a selection was created based on the stain and restored onto the APOE channel of the same brain section. Then APOE percent area in either astrocytes or neurons was measured.

For DG granule cell layer thickness measurements, two brain sections were stained according to the protocol above with anti-NeuN to highlight the cell layers. The thickness was measured manually in Fiji as follows: a line was drawn perpendicular to the NeuN cell layer at four locations in both sections, and an average was taken for each mouse.

### Western blot analysis

Biochemically extracted mouse neuroblastoma cell lysates were loaded onto 12% NuPAGE Bis-Tris polyacrylamide gel (Invitrogen, Cat. #NP0343) and separated by gel electrophoresis at 160 V using MOPS running buffer (Invitrogen, Cat. #NP0001). The separated proteins were transferred onto nitrocellulose membranes (Bio-Rad, Cat. #1620215) at 18 V for 60 min (Trans-Blot Turbo Transfer System, Bio-Rad). Membranes were washed for 3 × 5 min in PBS-T (PBS + 0.1% Tween-20; Millipore Sigma, Cat. #P2287) and then incubated in Intercept blocking buffer (LI-COR, Cat. #927-70001) for 1 h at room temperature to block nonspecific binding sites. After blocking, membranes were washed for 3 × 5 min in PBS-T and incubated with primary antibody overnight at 4 °C (APOE at 1:5000 dilution (Cell Signaling Technology, Cat. #13366) and beta actin at 1:2500 dilution (Abcam, Cat. #ab8226)). Membranes were washed for 3 × 5 min in PBS-T and incubated in fluorescently labeled secondary antibody: donkey anti-mouse 488 (1:2000; Invitrogen, Cat. #A21202) and donkey anti-rabbit 800RD (1:20000; LI-COR, Cat. #926-32213) for 1 h in the dark at room temperature. Resulting bands were detected with the ChemiDoc MP system (Bio-Rad), and the fluorescent intensity of each band was measured with Image Lab (Bio-Rad). The fluorescence intensity of the bands was quantified as a ratio of APOE:beta actin signal normalized to cells treated with ASO-NTC.

### Hippocampal volumetric analysis

Every tenth coronal brain section spanning the hippocampus was mounted on a microscope slide (Fisher Scientific, Cat. #22-037-246) for each mouse (seven sections per mouse, 30 μm thick, 300 μm apart) and dried for 1 hr at room temperature. Then, the brain sections were treated for 10 min with a 1% Sudan Black solution, washed with 70% ethanol for 2 min, washed with PBS-T for 2 min three times until the debris was washed away, and coverslipped with ProLong Gold mounting medium (Invitrogen, Cat. #P36930). To prepare the 1% Sudan Black solution, Sudan Black powder (Millipore Sigma, Cat. #S-2380) was combined with 70% ethanol (KOPTEC, Cat. #V1016) and mixed using a magnetic stirrer. The solution was then centrifuged at 1,100g for 10 min and the collected supernatant was filtered using a 0.2-μm filter syringe (Thermo Scientific, Cat. #723-2520). The stained, mounted brain sections were imaged on the Keyence microscope at x10 magnification. To quantify the volumes of the hippocampus and posterior lateral ventricle, we traced the areas of interest in ImageJ and used the formula: volume = distance between sections (30µm per section x 10 sections in between stained sample sections = 0.3mm) x sum of areas of all stained sections. We took a sum of seven brain sections per mouse, roughly between coordinates AP = −1.5 and AP = −3.3.

### Single-nuclei preparation for 10x loading

Single-nuclei preparation was conducted as previously described^11,25^. In brief, hippocampi previously dissected and stored at-80°C were combined with 1 ml of ice-cold homogenization buffer (HB) (250 mM sucrose, 25 mM KCl, 5 mM MgCl2, 20 mM tricine-KOH pH 7.8, 1 mM dithiothreitol, 0.5 mM spermidine, 0.15 mM spermine, 0.3% NP40) supplemented with 0.2 U/µl RNasin Plus RNAse inhibitor (Promega, Cat. #N2615) and protease inhibitor (Roche, Cat. 11836170001). Samples were dounced with ‘A’ loose pestle and ‘B’ tight pestle for 10 strokes each. Homogenate was filtered with a 70 µm Flowmi strainer (Bel-Art, Cat. #H13680-0070) and spun at 350 g for 5 min at 4°C. Supernatant was discarded and the pelleted nuclei were resuspended in 400 μL 1X HB. Next, 400 μl of 50% Iodixanol solution was added to the nuclei to create a 25% mixture, and then slowly layered with 600 μl of 30% Iodixanol solution underneath. Mixture was then layered again with 600 μl of 40% Iodixanol solution under the 30% mixture. The nuclei were spun at 3000 g for 20 min at 4°C before 200 μl of the nuclei band at the 30%–40%interface was combined with 800 μl of 2.5% BSA (Millipore Sigma; Cat. #A9576) in PBS supplemented with RNase inhibitor. Samples were spun for 10 min at 500 g and 4°C. The nuclei were resuspended with 2.5% BSA in PBS supplemented with RNase inhibitor to reach at least 500 nuclei per μl. Nuclei were then filtered with a 40 μm Flowmi stainer (Millipore Sigma; Cat. #BAH136800040) and counted. Roughly 14,000 nuclei per sample were loaded onto a 10x Genomics Next GEM chip G. The snRNAseq libraries were prepared using the Chromium Next GEM Single Cell 3ʹ Library and Gel Bead kit v.3.1 (10x Genomics, Cat. #1000121) according to the manufacturer’s instructions. Libraries were sequenced on an Illumina NovaSeq X Plus sequencer at the UCSF CAT Core.

### Pre-processing and clustering of mouse snRNA-seq samples

The snRNA-seq dataset included a total of 18 samples with 6 mice from each of the 3 treatment groups: PS19-E4 mice treated with ASO 4, PS19-E4 mice treated with ASO 75, and PS19-E4 mice treated with NTC ASO. The 6 PS19-E4 mice treated with ASO 4 had 4 male and 2 female mice; the 6 PS19-E4 mice treated with ASO 75 had 3 male and 3 female mice; and the 6 PS19-E4 mice treated with NTC ASO had 3 male and 3 female mice. The demultiplexed FASTQ files for these samples were aligned to a custom mouse reference genome using the 10x Genomics Cell Ranger Count pipeline^41^ (version 9.0.1), following the Cell Ranger documentation. To accommodate the Homo sapiens MAPT (NCBI reference sequence NM_001123066.4) and the H. sapiens APOE (NCBI reference sequence NM_001123066.4) genes, the custom mouse reference genome was made using the reference mouse genome sequence (GRCm38) from Ensembl (release 98) and the mouse gene annotation file from GENCODE (release M23). The H. sapiens MAPT sequence and H. sapiens APOE sequence were appended as separate chromosomes to the end of the mouse reference genome sequence and the corresponding gene annotations were appended to the filtered mouse reference gene annotation GTF file. The include-introns flag was set to TRUE to capture reads mapping to intronic regions.

The filtered count matrices generated by the Cell Ranger count pipeline for the 18 samples were processed using the R package Seurat^42^ (version 4.4.0) for single-nucleus analysis. Each sample was pre-processed as a Seurat object, then all 18 samples were merged into a single Seurat object. Low-quality nuclei were removed using the following criteria: fewer than 500 total UMI counts, fewer than 200 features, or a mitochondrial gene ratio higher than 0.25%. Sparse gene features expressed in fewer than 10 cells were also excluded. After quality control, normalization and variance stabilization were performed using SCTransform^43^ (version 0.4.2) method for initial parameter estimation. Graph-based clustering was conducted using the Seurat functions, FindNeighbors and FindClusters. Cells were embedded in a k-nearest neighbor (KNN) graph on the first 50 principal components (PCs) using Euclidean distance in the PCA space, and edge weights between two cells were refined using Jaccard similarity. Clustering via the Louvain algorithm (FindClusters) with 15 PCs and a resolution of 0.7 produced 38 distinct biologically relevant clusters, which were subsequently used for further analyses.

### Cell type assignment

Cell type identities were annotated following a method we previously described^44–46^. Briefly, Seurat’s AddModuleScore() function was applied to the object using features from lists of known brain marker genes (up to 8 per cell type)^45,47^. Further subdivisions of hippocampus cell types, such as CA1 and CA3 pyramidal cells, were identified using hippocampus-specific marker genes curated from Hipposeq (https://hipposeq.janelia.org). A module score of each cell type was computed for each individual cell, and each cell was assigned the cell type that produced the highest module score among all tested cell types. Cells with their top two scores within 10% of each other were classified as potential hybrids and excluded from downstream analyses. Cells with negative scores for all cell types computed were classified as negative and excluded from downstream analyses. The cell type assignments per cluster was evaluated by inspecting homogeneity, distribution, and spatial separation of clusters in UMAP space and the expression of major cell type marker genes across clusters. Clusters containing comparable proportions of two or more cell types were deemed unknown or mixed.

### Gene set enrichment analysis

DE genes between clusters of interest or between two genotype groups were identified using the FindMarkers Seurat function on the RNA assay data. This algorithm uses the Wilcoxon rank-sum test to identify DE genes between two populations. DE genes were limited to genes detected in at least 10% of the cells in either population and with at least 0.1 log_2_ fold change. Volcano plots with log_2_ fold change and p value from the DE gene lists were generated using the ggplot2 R package^48^ (version 3.5.2). The p values are based on a hypergeometric test and are adjusted for multiple testing using the Benjamini–Hochberg method^49^.

### Association between clusters and genotype

A GLMM_AM was implemented in the lme4 (version 1.1-37) R package^50^ and used to estimate the associations between cluster membership and the mouse model. These models were run separately for each cluster of cells. The GLM model was performed with the family argument set to the binomial probability distribution and with the ‘nAGQ’ parameter set to 10, corresponding to the number of points per axis for evaluating the adaptive Gauss–Hermite approximation for the log-likelihood estimation. Cluster membership of cells by sample was modeled as a response variable by a two-dimensional vector representing the number of cells from the given sample belonging to and not belonging to the cluster under consideration. The corresponding mouse ID from which the cell was derived was the random effect variable, and the animal model for this mouse ID was included as the fixed variable. The reference treatment was set to PS19-E4 mice treated with NTC ASO. The resulting p values for the estimated LOR across the three animal models (with respect to the PS19-E4 mice treated with NTC ASO) and clusters were adjusted for multiple testing using the Benjamini–Hochberg method^49^.

### Association between proportion of cell types and histopathological parameters

A GLMM_histopathology was implemented in the lme4 (version 1.1-37) R package^50^ and used to identify cell types whose proportions are significantly associated with changes in histopathology across the samples. These models were performed separately for each combination of all clusters and each histological parameter. The GLM model was performed with the family argument set to the binomial probability distribution family and with the ‘nAGQ’ parameter set to 1 corresponding to a Laplace approximation for the log-likelihood estimation. Cluster membership of cells by sample was modeled as a response variable by a two-dimensional vector representing the number of cells from the given sample belonging to and not belonging to the cluster under consideration. The corresponding mouse model from which the cell was derived was included as a random effect, and, further, the mouse ID within the given mouse model was modeled as a random effect as well. Note, this represents the hierarchical nature of these data for the GLMM, and the mouse models are first assumed to be sampled from a ‘universe’ of mouse models; this is then followed by sampling mice within each mouse model. The modeling choice of including the mouse model as a random effect as opposed to a fixed effect is meant to increase the degrees of freedom (or maximize the statistical power) to detect the association of interest, particularly in light of the relatively small number of replicates (3–4) per animal model. The histological parameter under consideration was modeled as a fixed effect in this model.

We visualized the LOR estimates (derived from the GLMM fits) in a heat map using pheatmap package 1.0.13 after adjusting the p values distribution across histopathological parameters across cell types with Benjamini–Hochberg multiple testing correction^49^.

### Subclustering of inhibitory neuronal, astrocytic, and microglial snRNA-seq data

The hippocampal cell clusters 8, 11, 15, and 26 were annotated as inhibitory neurons, cell cluster 13 was annotated as the astrocyte cells, and hippocampal cell clusters 6 and 21 were annotated as the microglial cells. Each of these cell types were further subclustered. Normalization and variance stabilization was performed using sctransform^51^ with the ‘glmGamPoi’ (Bioconductor package v.1.14.3) method^52^ for initial parameter estimation. Graph-based clustering was performed using the Seurat v.4.4.0 functions FindNeighbors and FindClusters. First, the cells were embedded in a k-nearest neighbor graph based on the Euclidean distance in the principal-component analysis space. The edge weights between two cells were further modified using Jaccard similarity. Next, clustering was performed using the Louvain algorithm implementation in the FindClusters Seurat function. Clustering was performed for all combinations of 10, 15, 20, 25 and 30 PCs with 0.4, 0.5, 0.6, 0.7, 0.8 and 0.9 resolutions. Subclustering with 15 PCs and 0.9 resolution resulted in 23 distinct biologically relevant subclusters for inhibitory neurons. Subclustering with 15 PCs and 0.9 resolution resulted in 15 distinct biologically relevant subclusters for astrocytes. Subclustering with 15 PCs and 0.9 resolution resulted in 13 distinct biologically relevant microglia subclusters.

### General Statistics and Reproducibility

Sample sizes for mouse studies were chosen on the basis of estimates to provide statistical power of ≥80% and alpha of 0.05 based on preliminary data. The differences between ASO treatment groups were evaluated by ordinary one-way ANOVA with Tukey’s multiple comparisons test, where the mean of each column was compared to the mean of every other column, or Welch’s ANOVA followed by Dunnett’s T3 multiple comparison test, where the mean of each column was compared to the mean of the control column. All data are shown as mean ± s.d. or ± s.e.m. Data distribution was assumed to be normal, though this was not formally tested. For correlations between two data sets, simple linear regression was used. p < 0.05 was considered to be significant, and all significant p values were included in figures or noted in figure legends. Statistical significance analysis and plots were completed with GraphPad Prism 10 for Mac (GraphPad Software). No randomization method was used for the assignment of mice to study groups, and no animals or data points were excluded from these studies.

For mouse snRNA-seq studies, sample sizes were determined by a power analysis based on effect sizes from our previous studies and literature^11,25^. Mice selected for snRNA-seq consisted of both males and females that together represented average pathologies for all parameters quantified. Nuclei were isolated from six mice per genotype to ensure n ≥ 3 mice per group. Pathology data were correlated with snRNA-seq data. Investigators were not blinded during analysis of the snRNA-seq datasets, as sample metadata were needed to conduct any comparisons. Studies were all performed using one cohort of mice.

## Data availability

The H. sapiens MAPT sequence is available at https://www.ncbi.nlm.nih.gov/nuccore/NM_001123066. The reference mouse genome sequence (GRCm38) from Ensembl (release 98) is available at http://ftp.ensembl.org/pub/release-98/fasta/mus_musculus/dna/Musmusculus.GRCm38.dna.primary_assembly.fa.gz. The reference mouse gene annotation file from GENCODE (release M23) is available at http://ftp.ebi.ac.uk/pub/databases/gencode/Gencode_mouse/release_M23/gencode.vM23.primary_as sembly.annotation.gtf.gz. The snRNA-seq datasets generated during the study are available at the Gene Expression Omnibus under accession number (link).

Data associated with Figs. 5 and 6 and Extended Data Figs. 3–8 are also available in the Supplementary Information. Source data are provided with this paper.

## Code availability

The following packages/software were used either as dependencies to downloading or using packages mentioned in the Methods section or in creating the figures in this manuscript: Seurat v.4.4.0, CellRanger v.9.0.1, glm lme4 v.1.1-37, clusterProfiler v.4.14.6, ggplot2 v.3.5.2, pheatmap v.1.0.12, and Fiji v.2.1.0 (ImageJ). Western blot images were analyzed using Image Lab v.6.1.0. GraphPad Prism v.10.6.1 was used for some statistical analyses and graphing. All codes generated with custom R and shell scripting during this study are accessible via GitHub at (link).

## Acknowledgements

This work was supported by National Institutes of Health (NIH) grants R10AG071697, RF1AG076647, and P01AG073082 to Y. Huang. The funders had no role in study design, data collection and analysis, decision to publish or preparation of the manuscript. We thank the entire Huang laboratory and Gladstone Institute of Neurological Disease colleagues for invaluable discussions and input; E. Chow and the staff at the UCSF Center for Advanced Technology Core for support with RNA-sequencing; and T. Pak for editorial assistance.

## Author Contributions

O.Y. and Y. Huang collaboratively designed the experiments and wrote the manuscript. O.Y. performed the majority of studies, data collection, and analyses. L.Y. and Y.L. performed the snRNA-seq analyses and generated sequencing data plots. J.B. aided in tissue collection, processing, and immunohistochemical staining. M.J.K., D.S., and Z.P. assisted in some tissue processing, immunohistochemical staining, and microscopy. K.S., A.A., and Y.Hao isolated cell nuclei and prepared samples for snRNA-seq. C.E. assisted with tissue collection. Q.X. assisted with ASO construct design. C.E., S.D.L., and J.N. managed all mouse lines. Y.Huang supervised the project.

---

**Extended Data Fig. 1.**
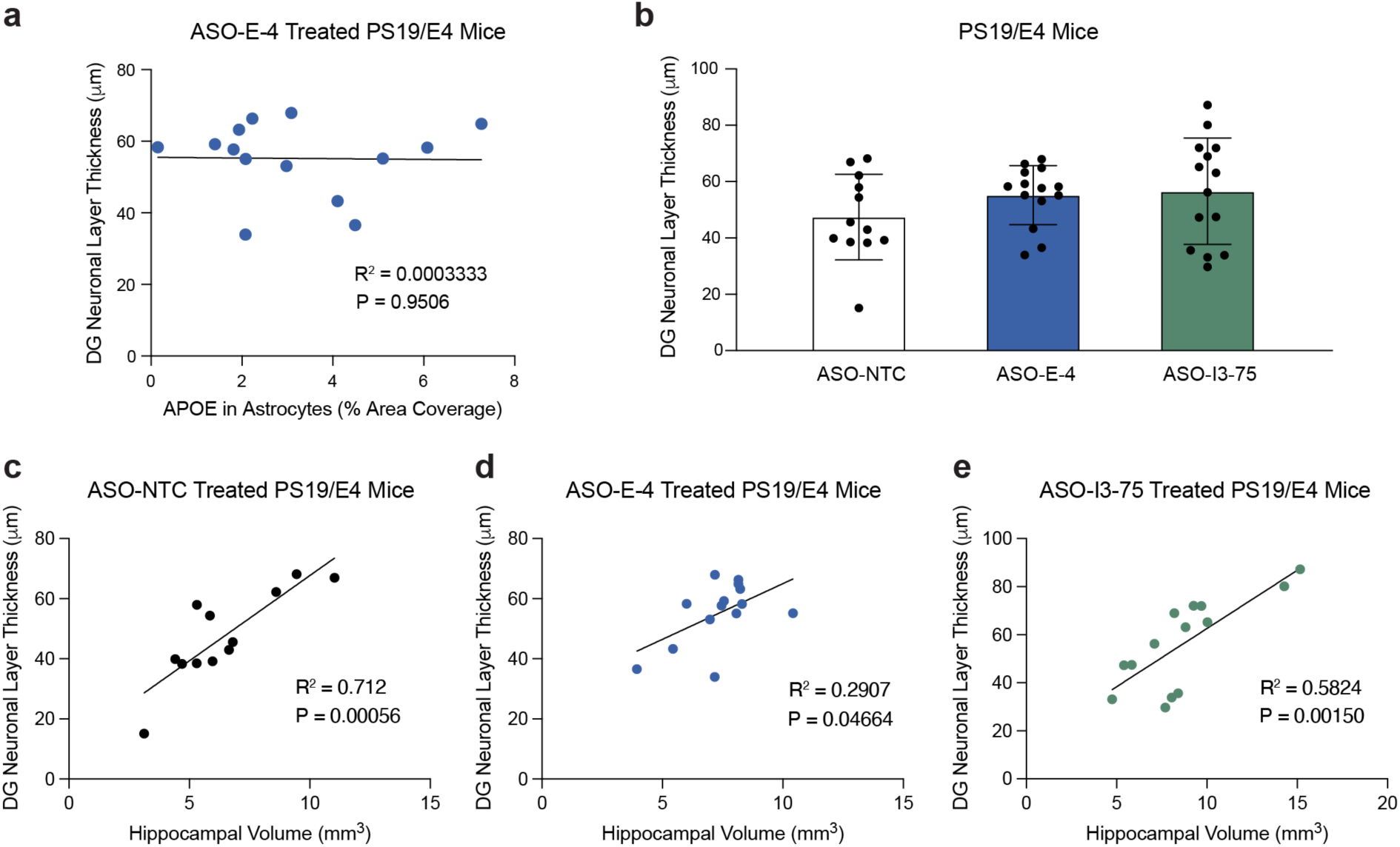
Trending Rescue of DG neuronal layer thickness in APOE ASO-treated PS19/E4 mice. a, Correlation between DG neuronal layer thickness (μm) and APOE in astrocytes (% area coverage) in PS19/E4 mice treated with ASO-E-4 (n = 14). **b**, Quantification of DG neuronal layer thickness (μm) in 9.5-month-old PS19/E4 mice treated with ASO-NTC, ASO-E-4, or ASO-I3-75 for 4 weeks. Data are expressed as mean ± s.d. and differences between groups were determined by ordinary one-way ANOVA followed by Tukey’s multiple comparison test. **c-e**, Correlations between DG neuronal layer thickness (μm) and hippocampal volume (mm^3^) in PS19/E4 mice treated with ASO-NTC (**c**, n=12), PS19/E4 mice treated with ASO-E-4 (**d**, n=14), and PS19/E4 mice treated with ASO-I3-75 (**e**, n=14). For all correlations, Pearson’s correlation analysis (two-sided) was performed.

**Extended Data Fig. 2.**
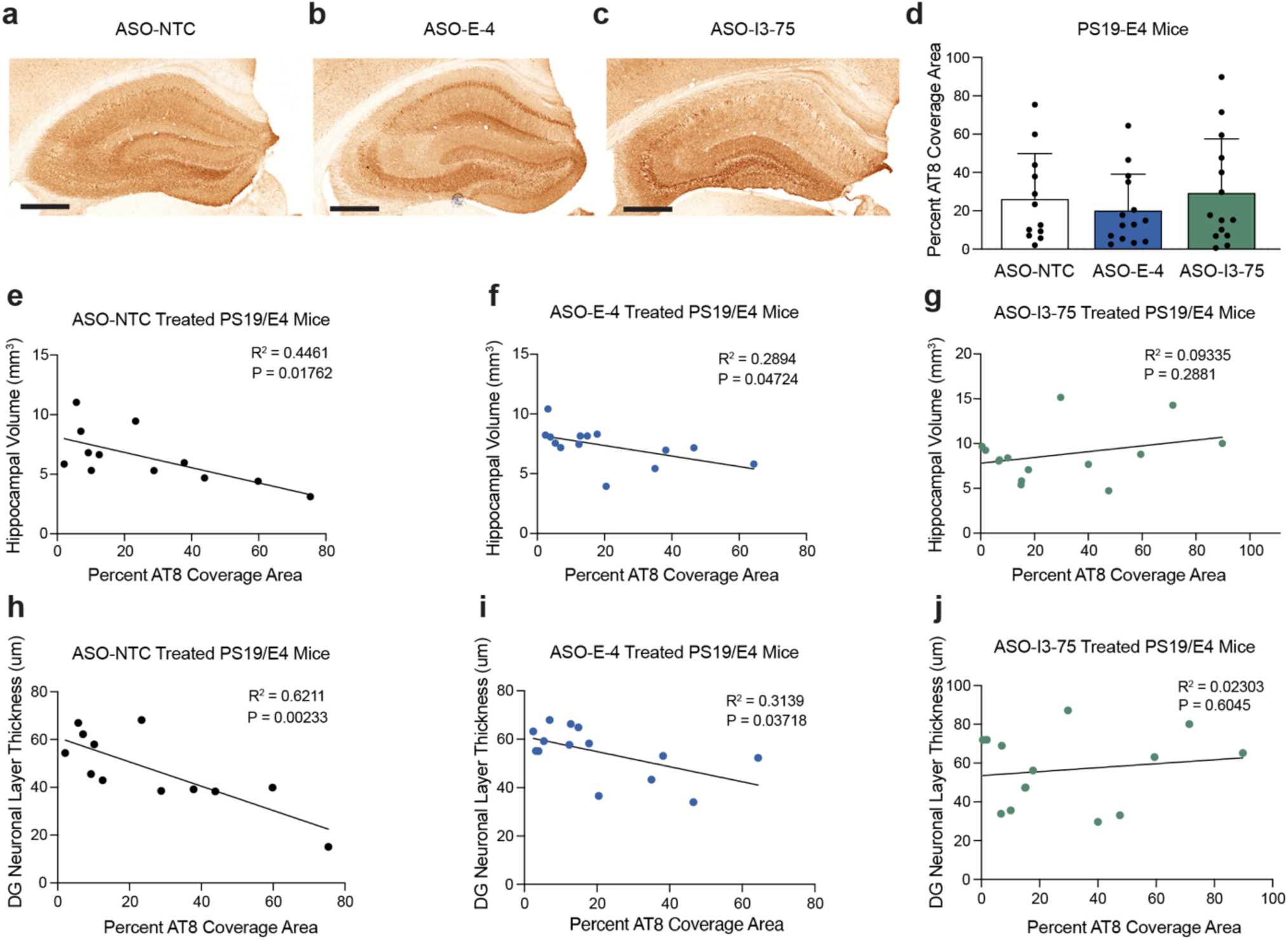
**ASO-I3-75, but not ASO-E-4, diminishes p-tau contribution to neurodegeneration in the hippocampus of PS19/E4 mice**. **a-c,** Representative images of p-tau immunostaining with the monoclonal antibody AT8 in the hippocampus of 9.5-month-old PS19/E4 mice treated with ASO-NTC (**a**), ASO-E-4 (**b**), or ASO-I3-75 (**c**) for 4 weeks (scale bar, 500μm). **d**, Quantifications of percent p-tau coverage area in the hippocampus of 9.5-month-old PS19/E4 mice treated with ASO-NTC, ASO-E-4, or ASO-I3-75 for 4 weeks. Data are expressed as mean ± s.d., and differences between groups were determined by ordinary one-way analysis of variance (ANOVA) with Tukey’s post hoc multiple comparisons test. **e-g**, Correlations between hippocampal volume (mm^3^) and percent AT8 coverage area in PS19/E4 mice treated with ASO-NTC (**e**, n=12), PS19/E4 mice treated with ASO-E-4 (**f,** n=14), and PS19/E4 mice treated with ASO-I3-75 (**g**, n=14). **h-j** Correlations between DG neuronal layer thickness (μm) and percent AT8 coverage area in PS19/E4 mice treated with ASO-NTC (**h**, n=12), PS19/E4 mice treated with ASO-E-4 (**i**, n=14), and PS19/E4 mice treated with ASO-I3-75 (**j**, n=14). For all correlations, Pearson’s correlation analysis (two-sided) was performed.

**Extended Data Fig. 3.**
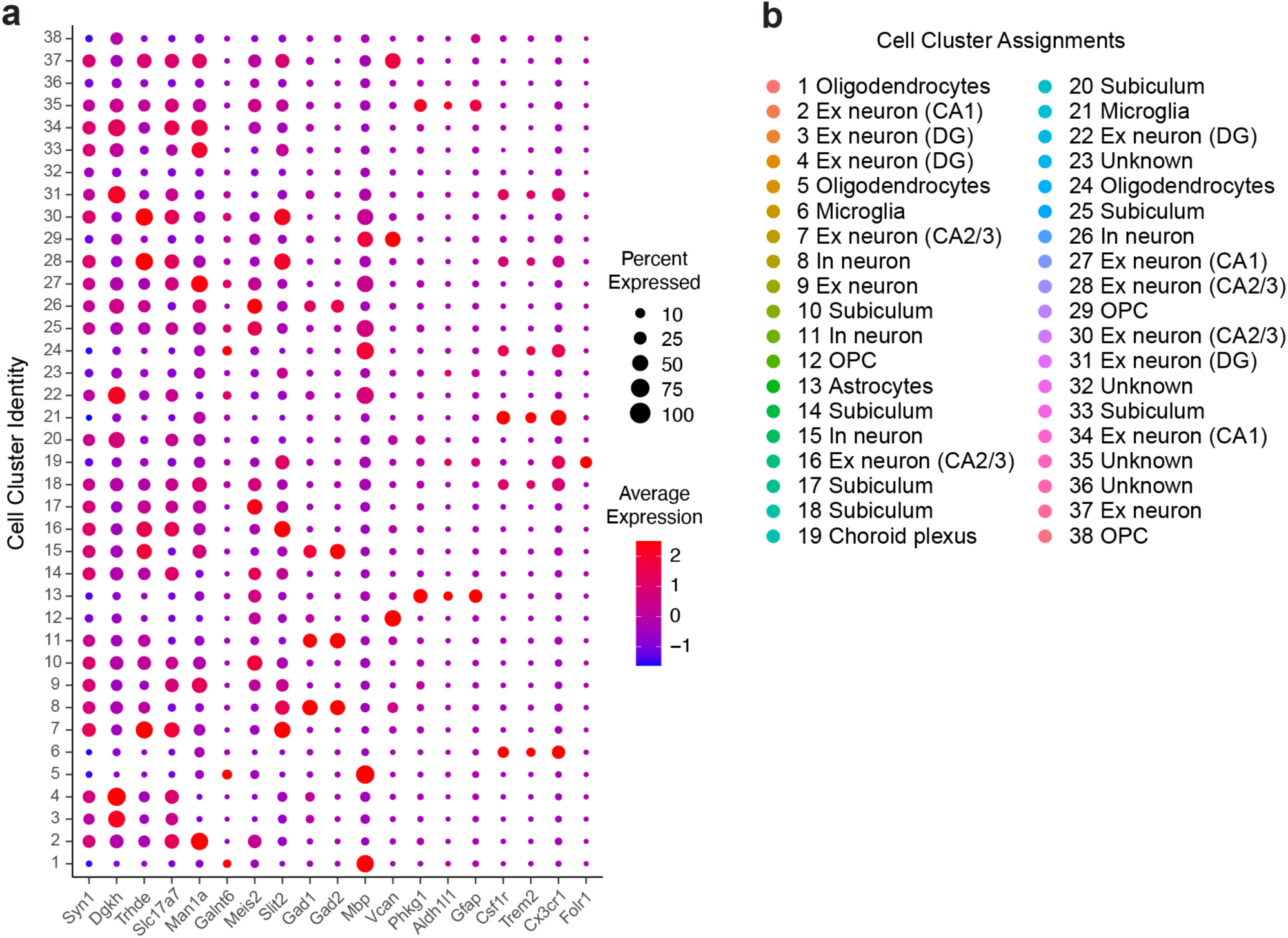
snRNA-seq identification of cell clusters. **a**, Dot-plot depicting normalized average expression of selected cell identity marker genes for all 38 unique hippocampal cell clusters from 9.5-month-old PS19/E4 mice treated with ASO-NTC, ASO-E-4, or ASO-I3-75 for 4 weeks. The size of the dots is proportional to the percentage of cells expressing a given gene. Average expression is on a colored scale (lower expression, blue; higher expression, red). **b**, Assigned identity of 38 distinct cell types. Ex neuron, excitatory neuron; In neuron, inhibitory neuron; DG, dentate gyrus; OPC, oligodendrocyte precursor cell.

**Extended Data Fig. 4.**
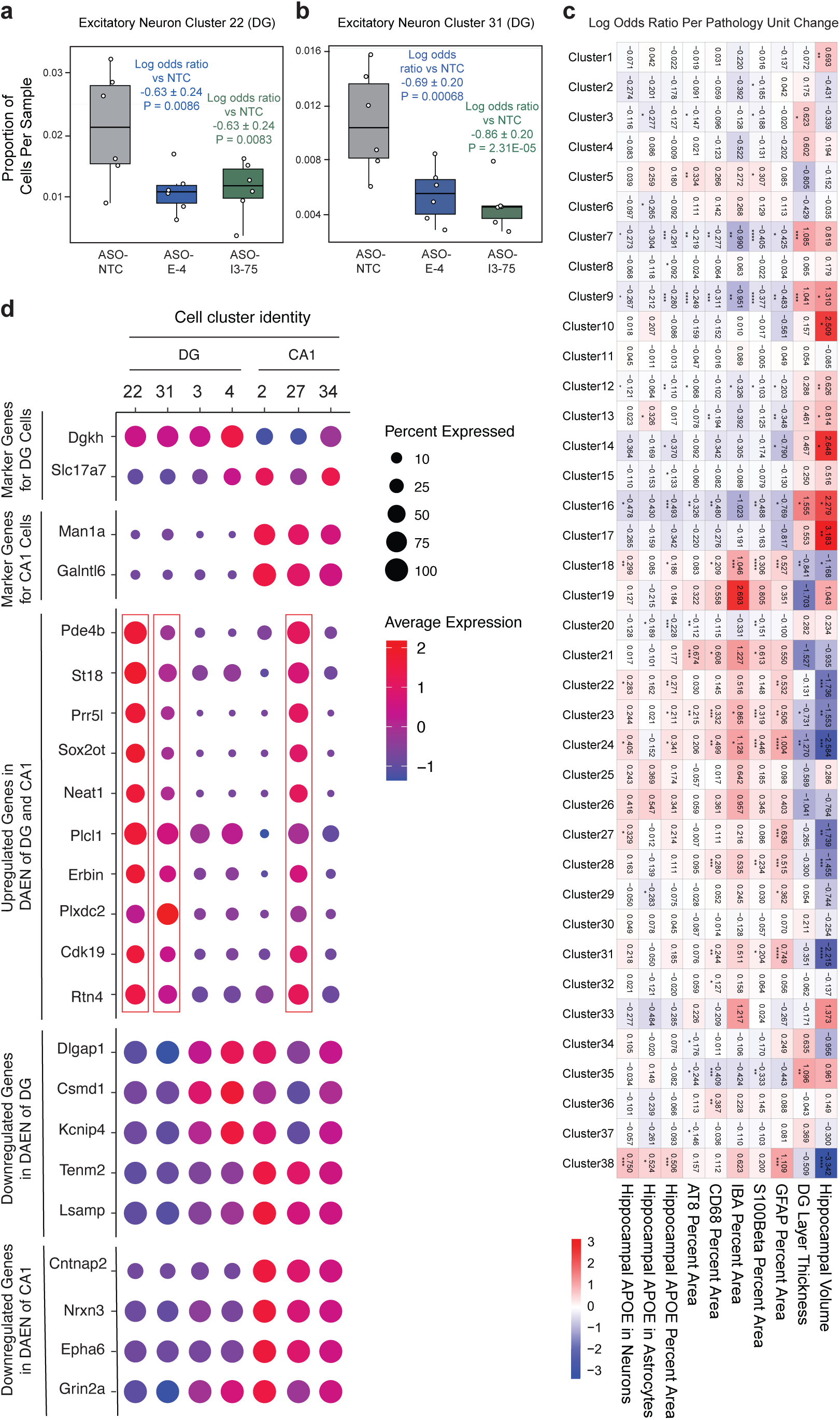
**ASO-E-4 and ASO-I3-75 diminishes disease-associated excitatory neurons (DAEN) in the hippocampus of ASO-treated PS19/E4 mice. a-b**, Box plot of the proportion of cells from each ASO treatment group in excitatory neuron cluster 22 (**a**) and excitatory neuron cluster 31 (**b**). For details see legend in Fig. 5b. **c**, Heat map plot of the log odds ratio per unit change in each pathological parameter for all 38 cell clusters. Negative associations are shown in blue and positive associations are shown in red. Unadjusted P values are from fits to a GLMM_ histopathology. **d**, Dot-plot of normalized average expression of marker genes and genes of interest for selected excitatory neuron clusters. The size of the dots is proportional to the percentage of cells expressing a given gene. Average expression is on a colored scale (lower expression, blue; higher expression, red).

**Extended Data Fig. 5.**
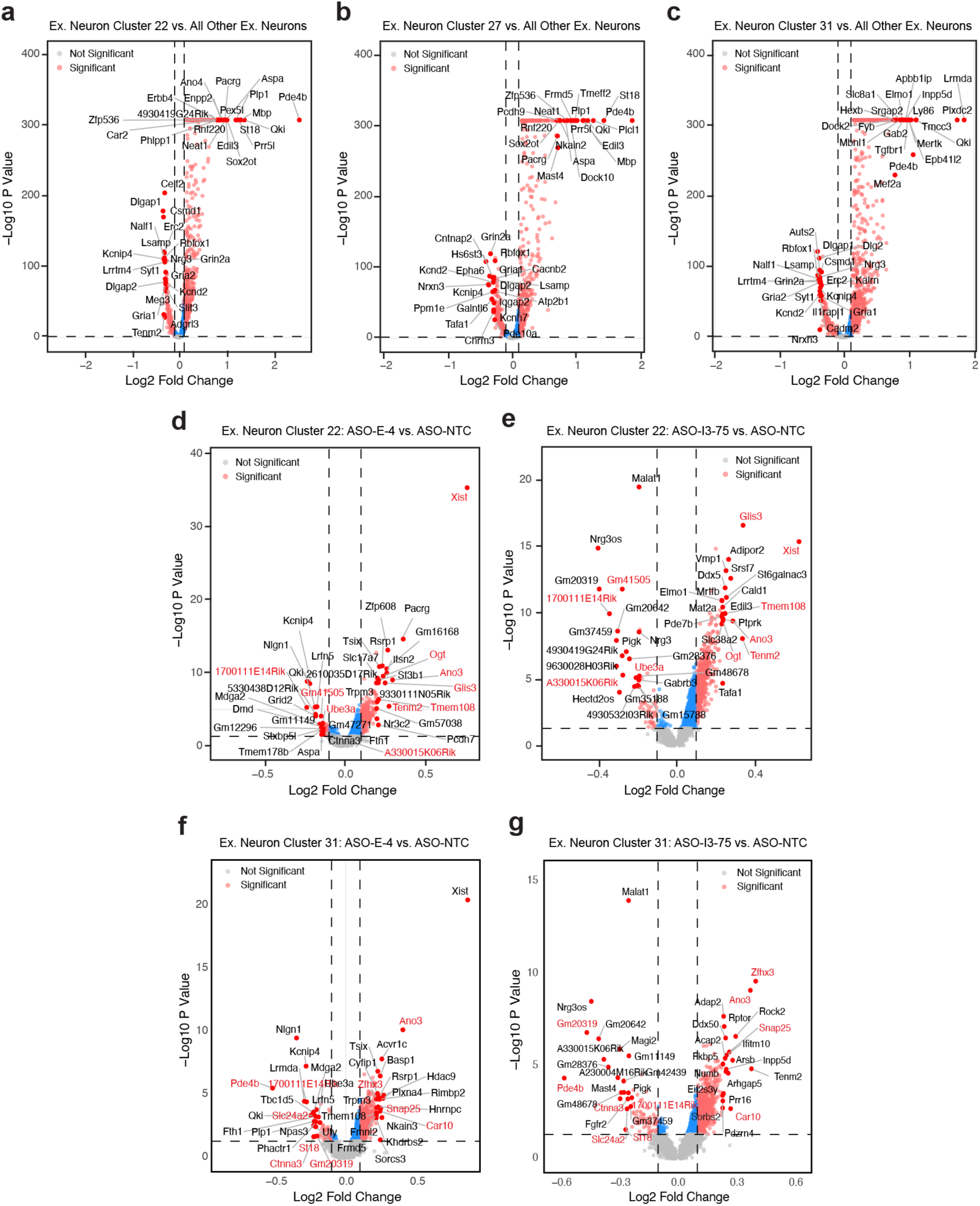
Differentially expressed genes (DEGs) in disease-associated excitatory neurons (DAEN) in the hippocampus of ASO-treated PS19/E4 mice. a-c, Volcano plot of the DEGs between excitatory neuron cluster 22 and all other excitatory neurons (**a**), between excitatory neuron cluster 27 and all other excitatory neurons (**b**), and between excitatory neuron cluster 31 and all other excitatory neurons (**c**). **d,e**, Volcano plot of the DEGs in excitatory neuron subcluster 22 between PS19/E4 mice treated with ASO-E-4 and those treated with ASO-NTC (**d**) and between PS19/E4 mice treated with ASO-I3-75 and those treated with ASO-NTC (**e**). Up- or down-regulated genes shared between **d** and **e** are indicated in red font. **f,g**, Volcano plot of the DEGs in excitatory neuron cluster 31 between PS19/E4 mice treated with ASO-E-4 and those treated with ASO-NTC (**f**) and between PS19/E4 mice treated with ASO-I3-75 and those treated with ASO-NTC (**g**). Up- or down-regulated genes shared between **f** and **g** are indicated in red font. For all volcano plots, dashed lines represent log_2_ fold change threshold of 0.1 and p value threshold of 0.05. The unadjusted p values and log_2_ fold change values used were generated from the gene-set enrichment analysis using the two-sided Wilcoxon rank-sum test as implemented in the FindMarkers function of the Seurat package.

**Extended Data Fig. 6.**
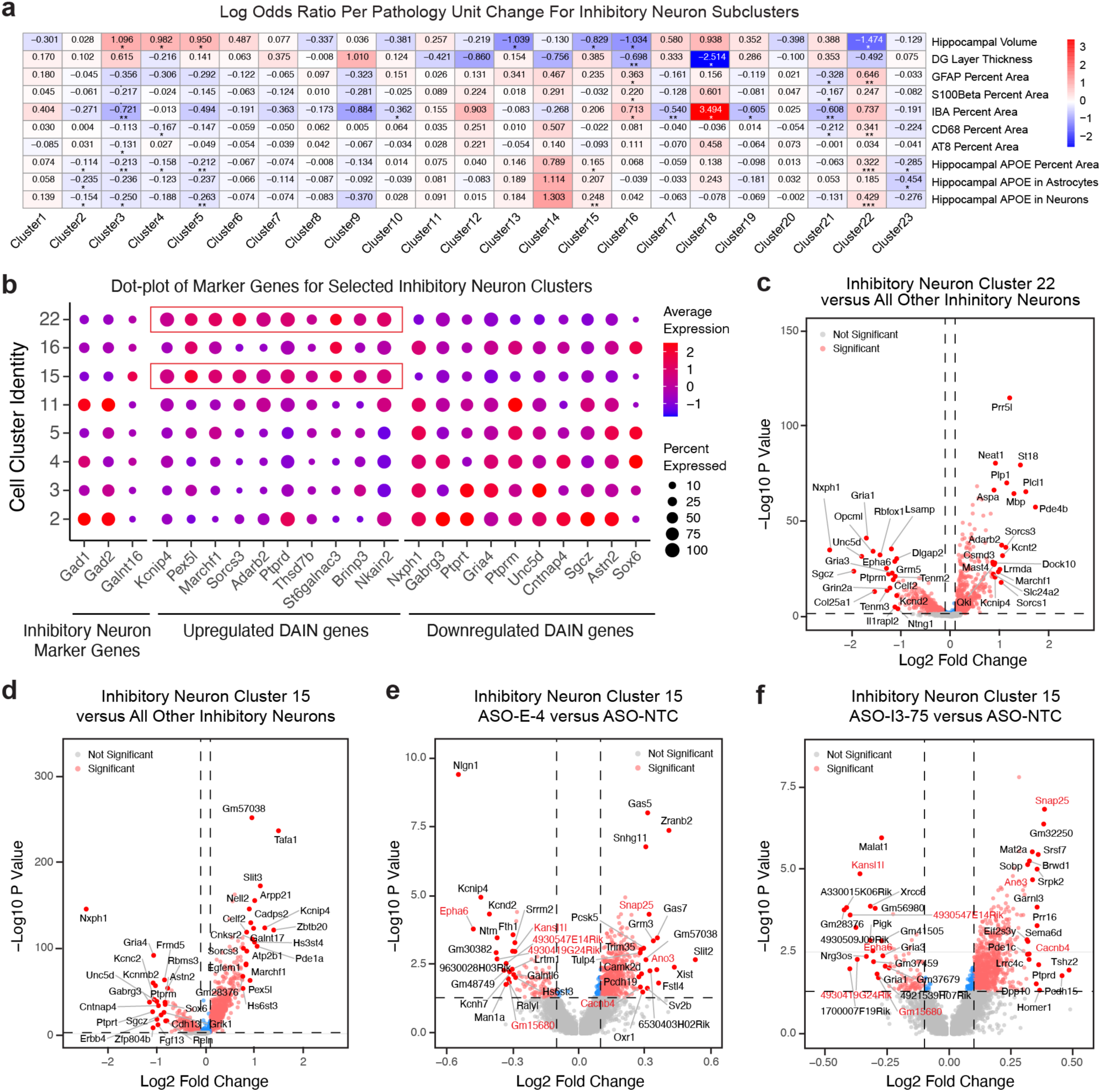
Characteristics of disease-associated inhibitory neurons (DAIN) in the hippocampus of ASO-treated PS19/E4 mice. a, Heat map plot of the log odds ratio per unit change in each pathological parameter for inhibitory neuron subclusters. Negative associations are shown in blue and positive associations are shown in red. Unadjusted P values are from fits to a GLMM_ histopathology. **b,** Dot-plot of normalized average expression of marker genes and genes of interest for selected inhibitory neurons. The size of the dots is proportional to the percentage of cells expressing a given gene. Average expression is on a colored scale (lower expression, blue; higher expression, red). **c,d**, Volcano plot of the DEGs between inhibitory neuron subcluster 22 and all other inhibitory neurons (**c**) and between inhibitory neuron subcluster 15 and all other inhibitory neuron (**d**). **e,f**, Volcano plot of the DEGs in inhibitory neuron subcluster 15 between PS19/E4 mice treated with ASO-E-4 and those treated with ASO-NTC **(e)** and between PS19/E4 mice treated with ASO-I3-75 and those treated with ASO-NTC (**f**). Up- or down-regulated genes shared between **e** and **f** are indicated in red font. For all volcano plots, dashed lines represent log2 fold change threshold of 0.1 and p value threshold of 0.05. The unadjusted p values and log2 fold change values used were generated from the gene-set enrichment analysis using the two-sided Wilcoxon rank-sum test as implemented in the FindMarkers function of the Seurat package.

**Extended Data Fig. 7.**
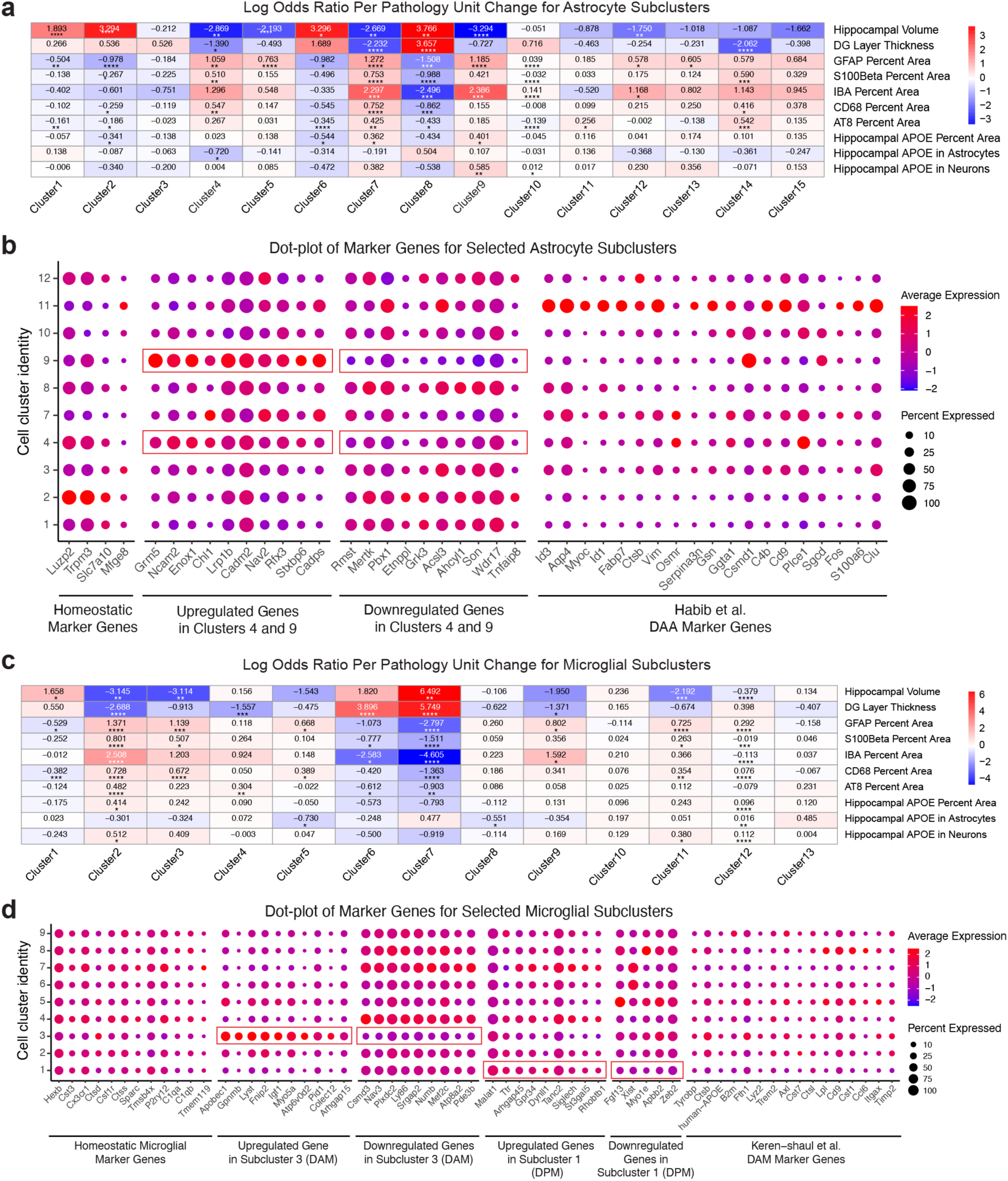
Characteristics of disease-associated astrocytes (DAA), disease-associated microglia (DAM), and disease-protective microglia (DPM) in the hippocampus of ASO-treated PS19/E4 mice. a, Heat map plot of the log odds ratio per unit change in each pathological parameter for all astrocyte subclusters. Negative associations are shown in blue and positive associations are shown in red. Unadjusted P values are from fits to a GLMM_ histopathology. **b,** Dot-plot of normalized average expression of marker genes and genes of interest for selected astrocyte subclusters. The size of the dots is proportional to the percentage of cells expressing a given gene. Average expression is on a colored scale (lower expression, blue; higher expression, red). **c**, Heat map plot of the log odds ratio per unit change in each pathological parameter for all microglia subclusters. Negative associations are shown in blue and positive associations are shown in red. Unadjusted P values are from fits to a GLMM_ histopathology. **d**, Dot-plot of normalized average expression of marker genes and genes of interest for selected microglia subclusters. The size of the dots is proportional to the percentage of cells expressing a given gene. Average expression is on a colored scale (lower expression, blue; higher expression, red).

**Extended Data Fig. 8.**
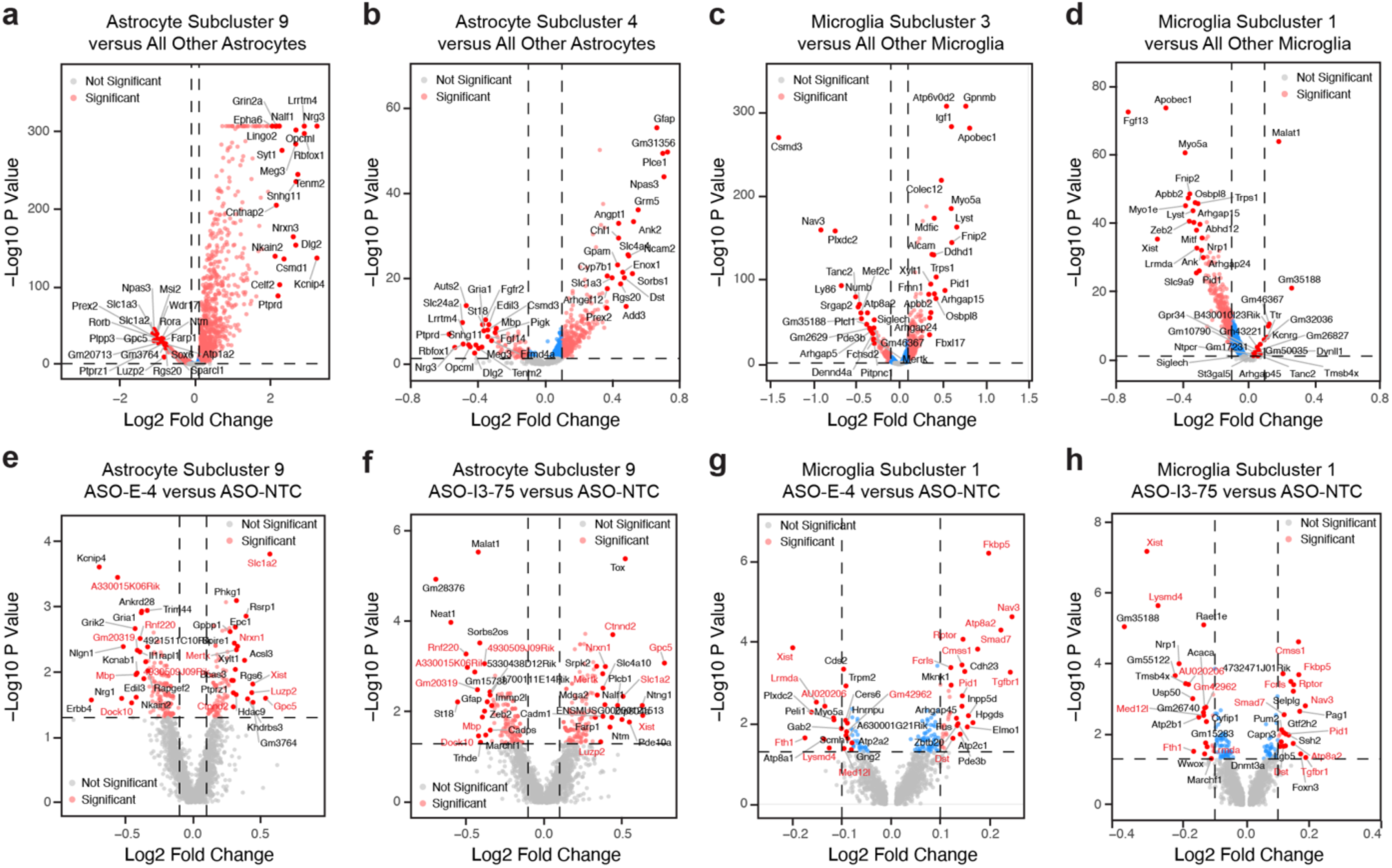
Differentially expressed genes (DEGs) in disease-associated astrocytes (DAA), disease-associated microglia (DAM), and disease-protective microglia (DPM) in the hippocampus of ASO-treated PS19/E4 mice. a,b, Volcano plot of the DEGs between astrocyte subcluster 9 and all astrocytes (**a**) and between astrocyte subcluster 4 and all other astrocytes (**b**). **c,d,** Volcano plot of the DEGs between microglia subcluster 3 and all other microglia (**c**) and between microglia subcluster 1 and all other microglia (**d**). **e,f**, Volcano plot of the DEGs in astrocyte subcluster 9 between PS19/E4 mice treated with ASO-E-4 and those treated with ASO-NTC **(e)** and between PS19/E4 mice treated with ASO-I3-75 and those treated with ASO-NTC (**f**). Up- or down-regulated genes shared between **e** and **f** are indicated in red font. **g,h**, Volcano plot of the DEGs in microglia subcluster 1 between PS19/E4 mice treated with ASO-E-4 and those treated with ASO-NTC **(g)** and between PS19/E4 mice treated with ASO-I3-75 and those treated with ASO-NTC (**h**). Up- or down-regulated genes shared between **g** and **h** are indicated in red font. For all volcano plots, dashed lines represent log2 fold change threshold of 0.1 and p value threshold of 0.05. The unadjusted p values and log2 fold change values used were generated from the gene-set enrichment analysis using the two-sided Wilcoxon rank-sum test as implemented in the FindMarkers function of the Seurat package.

